# Transcription-dependent regulation of replication dynamics modulates genome stability

**DOI:** 10.1101/286807

**Authors:** Marion Blin, Benoît Le Tallec, Viola Naehse, Mélanie Schmidt, Gael A. Millot, Marie-Noëlle Prioleau, Michelle Debatisse

**Affiliations:** Institut Curie, Centre de Recherche, Paris, France; Université Pierre & Marie Curie, Sorbonne Universités, Paris, France; CNRS UMR 3244, Paris, France; Institut Gustave Roussy, CNRS UMR 8200, Villejuif, France; Institut Pasteur, Bioinformatics and Biostatistics Hub, C3BI, USR 3756 IP CNRS, Paris, France; Institut Jacques Monod, CNRS UMR 7592, Université Paris Diderot, Paris, France; INSERM, Aix Marseille Université, CNRS, Institut Paoli-Calmettes, CRCM, Marseille, France; Ecole Normale Supérieure, Institut de Biologie de l’ENS (IBENS), INSERM U1024, CNRS UMR 8197, Paris, France; Department of Molecular Cell Biology, Institute for Cancer Research, The Norwegian Radium Hospital, Oslo, Norway

## Abstract

Replication stress is a primary threat to genome stability and has been implicated in tumorigenesis^1, 2^. Common fragile sites (CFSs) are loci hypersensitive to replication stress^3^ and are hotspots for chromosomal rearrangements in cancers^4^. CFSs replicate late in S-phase^3^, are cell-type dependent^4–6^ and nest within very large genes^4, 7–9^. The mechanisms responsible for CFS instability are still discussed, notably the relative impact of transcription-replication conflicts^7, 8, 10^ *versus* their low density in replication initiation events^5, 6^. Here we address the relationships between transcription, replication, gene size and instability by manipulating the transcription of three endogenous large genes, two in chicken and one in human cells. Remarkably, moderate transcription destabilises large genes whereas high transcription levels alleviate their instability. Replication dynamics analyses showed that transcription quantitatively shapes the replication program of large genes, setting both their initiation profile and their replication timing as well as regulating internal fork velocity. Noticeably, high transcription levels advance the replication time of large genes from late to mid S-phase, which most likely gives cells more time to complete replication before mitotic entry. Transcription can therefore contribute to maintaining the integrity of some difficult-to-replicate loci, challenging the dominant view that it is exclusively a threat to genome stability.

---

It is largely agreed that CFSs tend to remain incompletely replicated until mitosis upon replication stress. Incompletely replicated regions are processed by specific endonucleases promoting mitotic DNA synthesis and sister chromatid separation, eventually at the cost of chromosomal rearrangements^11–15^. Two main mechanisms have been suggested to explain this delayed replication completion. One postulates that secondary DNA structures^10^ or transcription-dependent replication barriers, notably R-loops^7, 8, 10^, lead to fork stalling and collapse. The other proposes that replication of the core of the CFSs by long-travelling forks due to their paucity in initiation events is specifically delayed upon fork slowing^5, 6^. Here we directly addressed the impact of transcription on the replication program and fragility of very large genes. For this purpose, we manipulated the transcription of such genes in chicken DT40 cells, where targeted DNA modification by homologous recombination is very efficient, and in human colon carcinoma HCT116 cells using the CRISPR-Cas9 technique.

We first focused on *DMD*, the largest annotated avian gene (Fig. 1). *DMD* extends over 996 kb and is neither transcribed (Fig. 1a) nor fragile (Fig. 1b) in DT40 cells. Determination of the replication timing showed that *DMD* replicates late (S4 fraction) in S-phase (Fig. 1e). We activated its transcription by inserting a tetracyclin-inducible promoter (Tet-promoter) on both alleles (Supplementary Fig. 1). A low level of *DMD* transcription from the Tet-promoter was detected in DMD^Tet/Tet^ cells grown without tetracycline (Fig. 1a). Importantly, fluorescent *in situ* hybridization (FISH) analysis of breaks induced in *DMD* by the DNA polymerase inhibitor aphidicolin revealed that the gene became fragile in these cells (Fig. 1b). Addition of tetracycline increased both *DMD* mRNA levels (Fig. 1a) and *DMD* instability (Fig. 1b). Our results therefore strongly support a major role of transcription in fragility, further illustrating that transcription constitutes an intrinsic source of genomic instability^16–18^.

**Figure 1.**
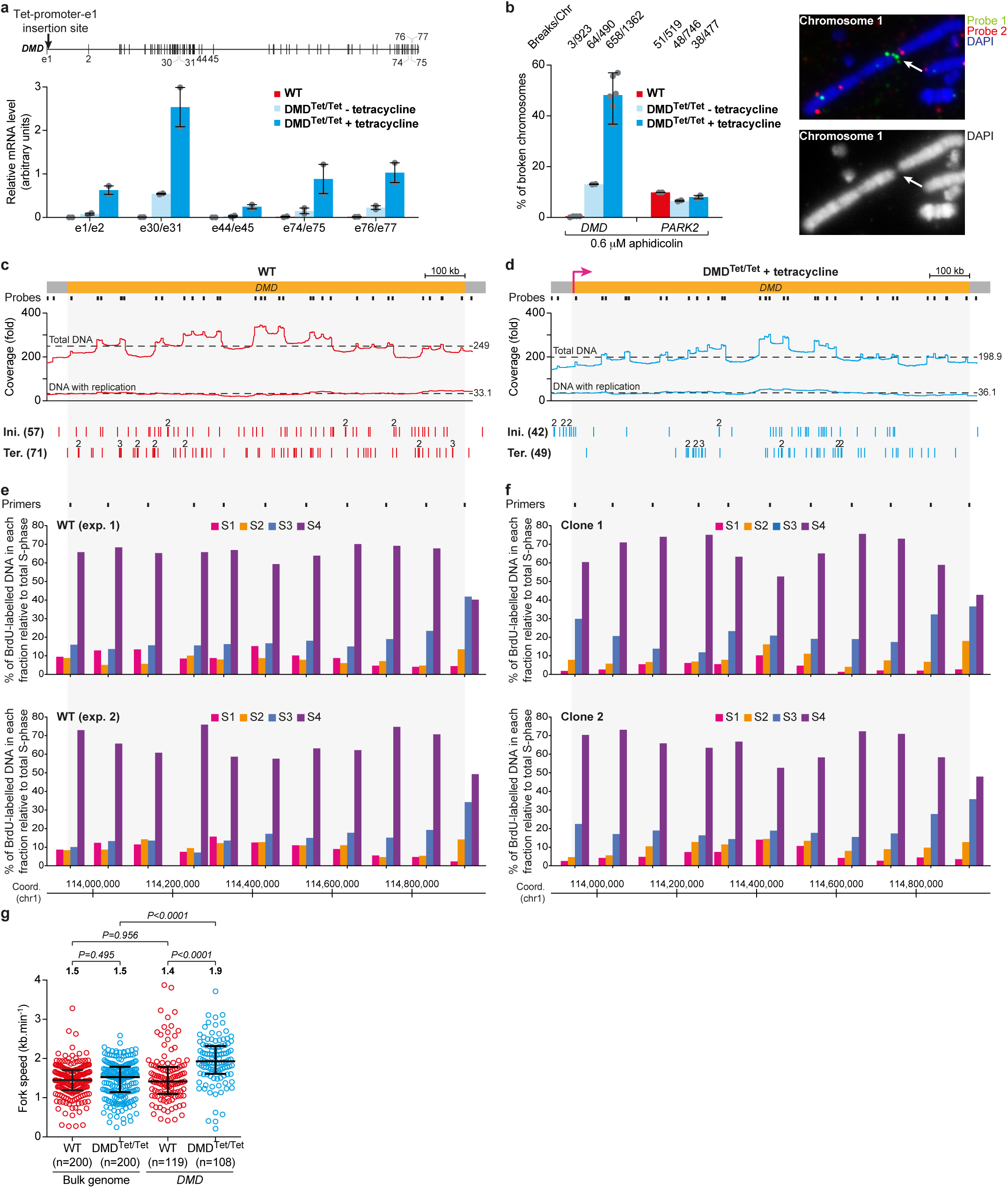
Impact of *DMD* transcription activation on its replication and fragility. **a**, *DMD* mRNA levels relative to β-actin mRNA in WT and DMD^Tet/Tet^ cells (median, extreme values and individual data points). Tested exonic junctions are indicated. Top: Map of *DMD* with its exons and the position of the Tet-promoter-*DMD* exon 1 insertion site. **b**, Left panel: Aphidicolin-induced breaks at *DMD* and *PARK2* in WT and DMD^Tet/Tet^ cells (median, extreme values and individual data points; aggregate numbers are presented on top). Breaks at the 679.5 kb-long *PARK2* gene were used as a control. Right panel: Example of two-color FISH with probes flanking *DMD*. Reverse-DAPI staining is also shown. The arrow indicates the position of a break. We also verified that *DMD* is not fragile in the absence of aphidicolin in WT and DMD^Tet/Tet^ cells (see Supplementary Fig. 1i). **c, d,** Mapping of initiation and termination events along the *DMD* locus in WT cells (**c**) and DMD^Tet/Tet^ cells with tetracycline (**d**). From top to bottom: *DMD* locus (yellow box. A pink arrow represents the active Tet-promoter in DMD^Tet/Tet^ cells) with the name of the cell line indicated above; Morse-code probes (black boxes) used in DNA-combing experiments to identify *DMD*; Total DNA and DNA with replication coverages of the locus (local peaks in total DNA coverage map to Morse-code probes; see Supplementary Figs 2, 3. Dotted lines represent mean coverages, indicated on the right); Map of initiation and termination events (the total number of events is indicated on the left. Figures above some lines indicate colocalized events). **e, f,** Replication timing profile of *DMD* in WT cells (**e**) and DMD^Tet/Tet^ cells with tetracycline (**f**). BrdU pulse-labelled cells were sorted into four S-phase fractions and neo-synthesized DNA was quantified by real-time PCR using the indicated primers (black boxes). Two experiments are shown; for DMD^Tet/Tet^ cells, replication timing analyses performed on two different clonal cell lines are presented. **g**, Replication fork progression in WT and DMD^Tet/Tet^ cells with tetracycline in the bulk genome or in *DMD*. Median with interquartile range (horizontal black lines), P-value and number of forks measured (n) are indicated.

We then examined the replication program of *DMD* in wild-type (WT) cells and in DMD^Tet/Tet^ cells grown in the presence of tetracycline. We used DNA-combing to visualize neo-synthesized DNA along single molecules spanning *DMD*, which permitted us to measure the speed of individual forks and to map initiation and termination events along the locus^5^ (Supplementary Figs 2, 3a and Supplementary Table 1). Initiation and termination events appeared randomly distributed along *DMD* in WT cells (Fig. 1c). In striking contrast, initiation events accumulated in the Tet-promoter region when *DMD* was transcribed (Fig. 1d). This initiation zone was followed by a large region depleted of initiation events extending over the 5’ part of the gene. Initiation events also clustered in a second, ≈300 kb-large zone located in the 3’ half of active *DMD.* As expected, modifications of the initiation profile resulted in a redistribution of termination events. Active *DMD* remained late-replicating (Fig. 1f), confirming that transcription *per se* is not sufficient to impose early replication^19, 20^. Thus, our results show that transcription modulates origin distribution, in line with recent analyses^21–23^, and account for the frequent localization of origins near transcription start sites^24, 25^. In addition, activation of *DMD* transcription specifically increased fork speed along the *DMD* locus (Fig. 1g). One hypothesis is that transcription could regulate chromatin permissiveness to fork progression. Consistently, chromatin factors like the histone chaperone FACT (Facilitates Chromatin Transcription), which participates in the disassembly of the nucleosomes upstream of the RNA polymerase, influence the rate of fork progression on *in vitro* reconstituted chromatin templates^26^.

**Figure 2.**
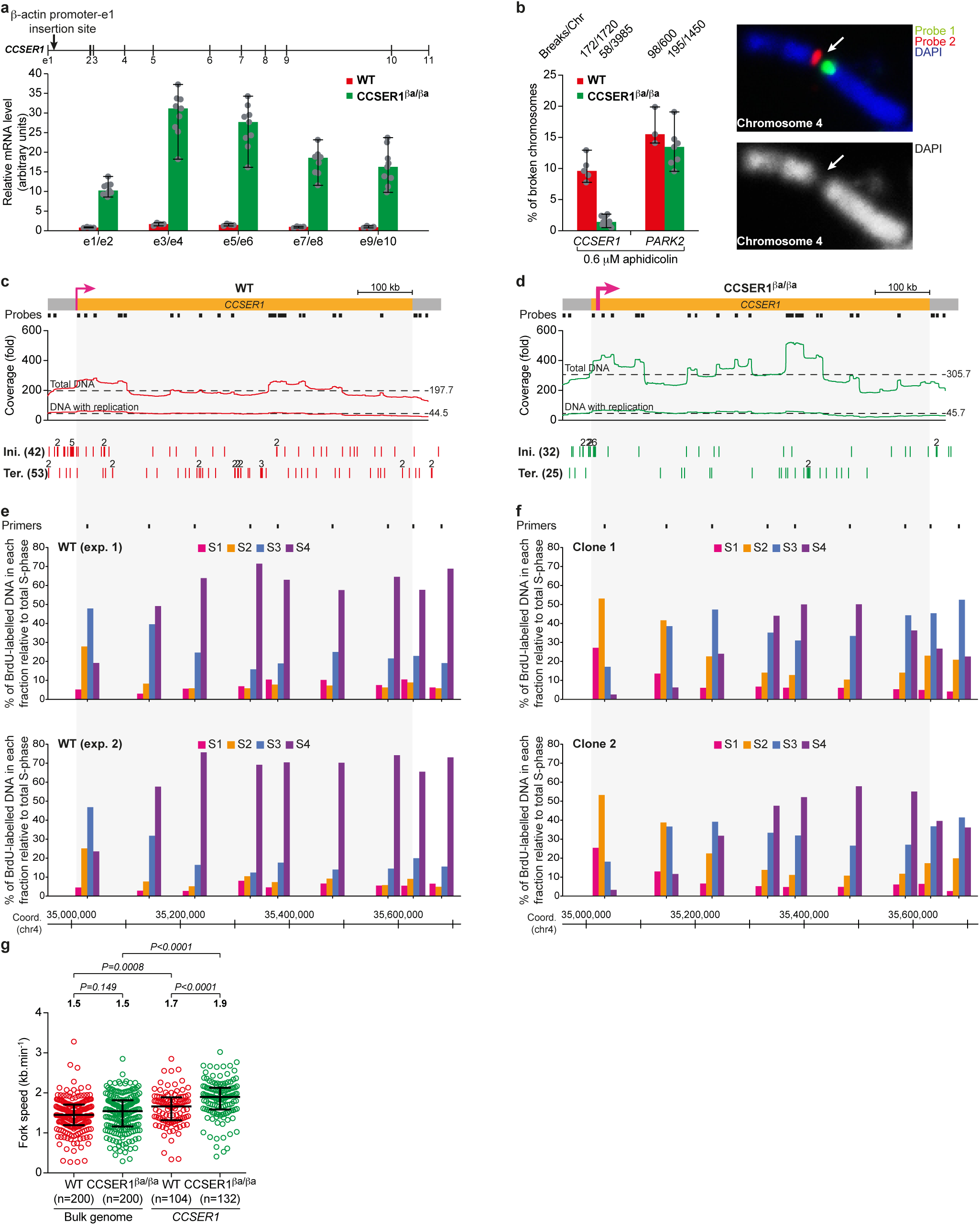
Impact of *CCSER1* transcription upregulation on its replication and fragility. **a**, *CCSER1* mRNA levels relative to β-actin mRNA in WT and CCSER1^βa/βa^ cells (median, extreme values and individual data points). Tested exonic junctions are indicated. Top: Map of *CCSER1* with its exons and the position of the chicken β-actin promoter-*CCSER1* exon 1 insertion site. **b**, Left panel: Aphidicolin-induced breaks at *CCSER1* and *PARK2* in WT and CCSER1^βa/βa^ cells (median, extreme values and individual data points; aggregate numbers are presented on top). Breaks at *PARK2* were used as a control. Right panel: Example of two-color FISH with probes flanking *CCSER1*. See Fig. 1b for details. We also verified that *CCSER1* is not fragile in the absence of aphidicolin in WT and CCSER1^βa/βa^ cells (see Supplementary Fig. 4g). **c, d,** Mapping of initiation and termination events along the *CCSER1* locus in WT (**c**) and CCSER1^βa/βa^ (**d**) cells. From top to bottom: *CCSER1* locus (yellow box; a thin and a thick pink arrow represents active *CCSER1* promoter in WT cells and active β-actin promoter in CCSER1^βa/βa^ cells, respectively) with the name of the cell line indicated above; Morse-code probes (black boxes) used in DNA-combing experiments to identify *CCSER1*; Coverage of the locus; Map of initiation and termination events. See Fig. 1c for details. **e, f,** Replication timing profile of *CCSER1* in WT (**e**) and CCSER1^βa/βa^ (**f**) cells. Two experiments are shown; for CCSER1^βa/βa^ cells, replication timing analyses performed on two different clonal cell lines are presented. See Fig. 1e for details. **g**, Replication fork progression in WT and CCSER1^βa/βa^ cells in the bulk genome or in *CCSER1*. See Fig. 1g for details.

**Figure 3.**
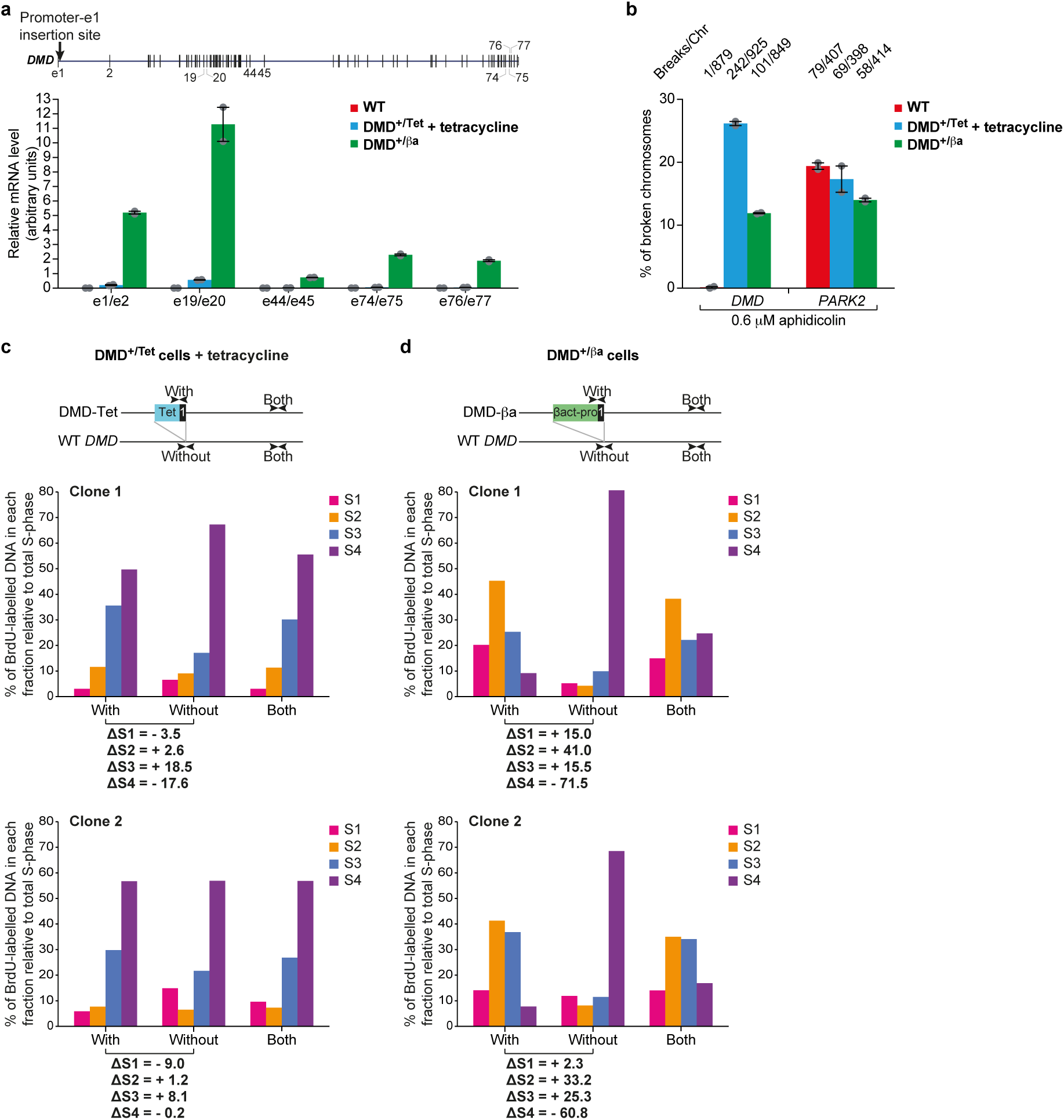
Impact of *DMD* transcription modulation on its replication timing and fragility. **a**, *DMD* mRNA levels relative to β-actin mRNA in WT cells, DMD^+/Tet^ cells with tetracycline and DMD^+/βa^ cells (median, extreme values and individual data points). Tested exonic junctions are indicated. Top: Map of *DMD* with its exons and the position of the Tet- or β-actin promoter-*DMD* exon 1 insertion site. **b**, Aphidicolin-induced breaks at *DMD* and *PARK2* in WT cells, DMD^+/Tet^ cells with tetracycline and DMD^+/βa^ cells (median, extreme values and individual data points; aggregate numbers are presented on top). Breaks at *PARK2* were used as a control. **c, d,** Replication timing at the Tet-(**c**) and β-actin (**d**) promoter insertion site. BrdU pulse-labelled DMD^+/Tet^ cells with tetracycline and DMD^+/βa^ cells were sorted into four S-phase fractions, and neo-synthesized DNA was quantified by real-time PCR using primer pairs specific of the DMD-Tet or DMD-βa allele (with), of the WT *DMD* allele (without) or hybridizing on the two alleles (both). The differences in replication timing at the target site following promoter integration (ΔS=S_with_-S_without_) were calculated for each sub-fraction. Results for two different clonal cell lines are presented for each construct.

We next modulated the transcription of the 616 kb-long *CCSER1* gene in DT40 cells (Fig. 2). *CCSER1* is transcribed (Fig. 2a) and is one of the most fragile regions of the DT40 genome upon aphidicolin treatment (Fig. 2b and ^4^). We enhanced *CCSER1* transcription by placing both alleles of the gene under the control of the strong chicken β-actin promoter (Supplementary Fig. 4). Unexpectedly, the ensuing ≈20 fold increase in *CCSER1* transcription (Fig. 2a and Supplementary Fig. 4f) was accompanied by a dramatic reduction of aphidicolin-induced *CCSER1* instability (Fig. 2b). To understand why, we compared *CCSER1* replication in WT and CCSER1^βa/βa^ cells (Fig. 2c-g, Supplementary Fig. 3b and Supplementary Table 1). A strong initiation zone overlapped *CCSER1* promoter region in WT cells, with some additional initiation events apparently randomly distributed along the gene (Fig. 2c).

Importantly, *CCSER1* overexpression further favoured initiation in the promoter region at the expense of the rest of the gene (Fig. 2d). In addition, termination events clustered in the third quarter of *CCSER1* in CCSER1^βa/βa^ cells (Fig. 2d), suggesting that CCSER1^βa^ allele was mainly replicated by forks proceeding inward from the initiation zone located upstream of the β−actin promoter and from a second initiation region located 3’ of *CCSER1*. Together with *DMD*, our results therefore demonstrate that transcription quantitatively dictates the replication initiation program of active large genes, as well as the ensuing termination profile. In agreement with the results obtained for *DMD*, enhancing *CCSER1* transcription also significantly increased fork velocity inside the locus (Fig. 2g).

Remarkably, *CCSER1* overexpression advanced the replication time of the 5’ part of the gene, which shifted from mid (S2/S3 fractions) to early S-phase (S1/S2 fractions) (Fig. 2f). It also markedly advanced the replication time of the 3’ end of *CCSER1* and its flanking region (Fig. 2f). These data are consistent with those obtained by combing (Fig. 2d), reciprocally validating each other and further supporting that transcription, at least at a sufficient level, stimulates initiation both upstream and downstream of large genes. One intriguing possibility is that the coordinated timing shift observed in 5’ and 3’ of *CCSER1* upon overexpression might be mediated by long-range chromatin interactions. Advanced initiation at both ends of *CCSER1* collectively led to a replication timing shift of the whole *CCSER1* locus in CCSER1^βa/βa^ cells, mainly from late (S4 fraction) to mid-late (S3/S4 fractions) S-phase (Fig. 2e, f). These results suggest that the stability of CCSER1^βa^ alleles stems from the advancement of replication timing induced by the strong increase in transcription, which gives cells more time to complete replication before entering mitosis. Together with the data obtained for *DMD*, they also suggest that only high transcription levels may be capable of advancing the replication timing of large genes.

To test this hypothesis, we used the β-actin promoter in place of the Tet-promoter to increase *DMD* transcription (Supplementary Fig. 5a-f). We failed to recover homozygous cells but heterozygous DMD^+/βa^ cells were alive and exhibited no growth defect (Supplementary Fig. 5g). We therefore compared these cells to heterozygous DMD^+/Tet^ cells treated with tetracycline (Fig. 3). *DMD* transcription was ≈10 times higher in DMD^+/βa^ than in DMD^+/Tet^ cells (Fig. 3a and Supplementary Fig. 5h) yet, in line with *CCSER1* results, *DMD* was two-fold less fragile upon aphidicolin exposure (Fig. 3b). As anticipated, we detected a strong shift to earlier replication timing at the β-actin promoter insertion site, from S4 to S2/S3 fractions (Fig. 3c, d).

Finally, we sought to extend our conclusions to FRA3B, the archetypal human CFS nested in the 1.5-Mb-long *FHIT* tumour suppressor gene^27^ (Fig. 4). We used the CRISPR-Cas9 system to up-regulate *FHIT* transcription in HCT116 epithelial cells, where FRA3B is the most active CFS^4^ (Supplementary Fig. 6). *FHIT* promoter region was substituted on both alleles with a cassette containing the strong human EF1α promoter (Supplementary Fig. 6), which resulted in a ≈20 fold increase in *FHIT* mRNA levels (Fig. 4a). This increase elicited a massive reduction of FRA3B instability upon aphidicolin treatment (Fig. 4b) accompanied by a shift to earlier replication of the *FHIT* locus, from late (S3/S4 fractions) to mid S-phase (S2/S3 fractions) (Fig. 4c, d). Thus, data from human cells are perfectly in agreement with those from DT40 cells, further substantiating that transcription directly and quantitatively influences the replication time of large genes and that advancing replication timing protects active large genes from fragility. Interestingly, the 5’ and 3’ ends of *FHIT* replicated earlier than its central part in WT cells (Fig. 4c). This profile is accentuated in FHIT^EF1α/EF1α^ cells (Fig. 4d), reminiscent of the replication timing profile of CCSER1^βa^ alleles (Fig. 2f). These results confirm that high levels of transcription favour advanced initiation of replication both 5’ and 3’ of active large genes, contributing to an earlier replication of the entire locus. Noticeably, it has been reported that the insertion in the genome of DT40 cells of the strong β-actin promoter upstream of the 423 bp-long blasticidin resistance gene has little impact on the replication timing of the targeted chromosomal region^19^, suggesting that transcription-dependent shift in timing may be limited to genes over a certain size.

**Figure 4.**
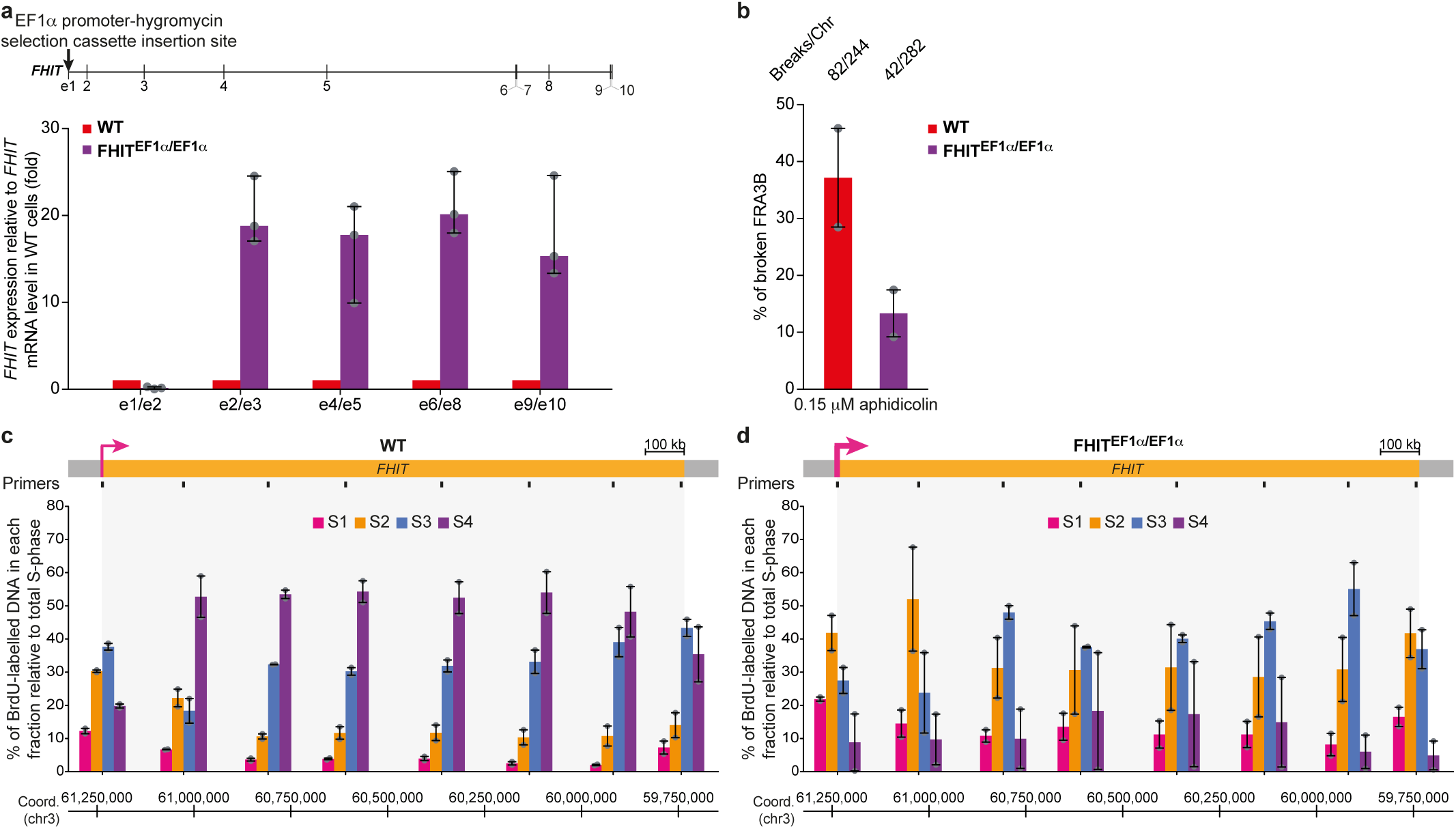
Impact of *FHIT* transcription upregulation on its replication timing and fragility in human HCT116 cells. **a**, *FHIT* expression in WT and FHIT^EF1α/EF1α^ cells relative to *FHIT* mRNA in WT cells (median, extreme values and individual data points). *FHIT* mRNA level was normalized to 3 housekeeping genes (*POLR2F*, *RPL11* and *PPIB*). Tested exonic junctions are indicated. Top: Map of *FHIT* with its exons and the position of the EF1α promoter-hygromycin selection cassette insertion site. **b**, Aphidicolin-induced breaks at *FHIT* in WT and FHIT^EF1α/EF1α^ cells (median, extreme values and individual data points; aggregate numbers are presented on top). **c, d,** Replication timing profile of *FHIT* in WT (**c**) and FHIT^EF1α/EF1α^ (**d**) cells (median and extreme values of 2 experiments). See Fig. 1e for details.

We show here that the degree of instability of large genes upon replication stress relies on their transcription level in an atypical way, with low transcription being more deleterious than high transcription levels. Strikingly, while cause-and-effect relationships between transcriptional regulation and replication timing have fuelled an intense debate^28^, we demonstrate that transcription regulates replication timing in a quantitative rather than qualitative manner. Noticeably, high levels of transcription advance the replication time from late to mid S-phase of all the three large genes that we studied and protect them from replication stress-induced breaks, which establishes a causal role for late replication timing in CFS instability. The high transcriptional-level dependent shift in replication timing most likely gives cells more time to complete replication prior to the onset of mitosis. Importantly, it was observed in multiple cell types that active large genes are generally transcribed at low levels^8, 9^ and, accordingly, tend to replicate late during S-phase^8^. Our results are therefore in line with recent genome-wide analyses concluding that CFSs specifically correspond to large, transcribed, late-replicating genes and also explain why not all transcribed large genes are fragile^4, 29^.

The role of transcription in origin distribution observed for both *DMD* and *CCSER1* prompted us to propose that initiation-poor regions and associated long–travelling forks observed in CFSs^3, 5, 6^ actually originate from and reflect the transcription of the cognate large genes. Whether CFS instability results from transcription-driven paucity in initiation events or transcription-dependent formation of barriers impeding fork progression or both phenomena remains to be determined.

In conclusion, while transcription is considered today as a primary threat to genome stability, our study reveals complex relationships between transcription, replication and chromosome fragility. The multifaceted impact of transcription on replication calls for a careful assessment of the mutational consequences of transcriptional changes, especially in cancer cells where up-regulation of transcription is suspected to be a major contributor of oncogene-induced replication stress^30^.

## Acknowledgments

We thank S. Lambert, O. Hyrien, F. De Carli, M. Hennion, L. Lacroix and V. Besic for critical reading of the manuscript. The authors would like to acknowledge the Cell and Tissue Imaging Platform - PICT-IBiSA (member of France–Bioimaging) of the Genetics and Developmental Biology Department (UMR3215/U934) of Institut Curie for help with light microscopy, the Flow Cytometry Platform Imagoseine of Institut Jacques Monod, Université Paris Diderot, and the Imaging and Cytometry Platform (PFIC) of Institut Gustave Roussy for assistance with cell sorting. M. D. team is supported by the Agence Nationale de la Recherche (ANR-13-BSV6-0008-01/FRA-Dom), the Association pour la Recherche sur le Cancer (Subvention Libre Sl220130607073) and the Institut National du Cancer (INCa subvention 2013-103). M. N. P. team is supported by the Association pour la Recherche sur le Cancer (Labellisation PGA120150202272) and the Agence Nationale de la Recherche (ANR-15-CE12-0004-01). M. B. was supported by fellowships from the Ministère de l’Enseignement Supérieur et de la Recherche and the Ligue contre le cancer.

## Methods

### Cell culture

DT40 cells were grown in RPMI 1640 GlutaMAX medium (Gibco) with 10% fetal bovine serum (Biowest), 2% chicken serum (Gibco) and 0.1 mM β-mercaptoethanol (Millipore). A tetracycline-free fetal bovine serum (Biowest) was used when appropriate. HCT116 human colon carcinoma cells were cultured in McCOY’S 5A medium (Gibco) with 10% fetal calf serum and 20 mM L-glutamine (Gibco) except for replication timing experiments for which they were cultured in DMEM (4.5 g/L D-Glucose, L-glutamine) (Gibco) with 10% fetal calf serum and 1 mM sodium pyruvate. All media were supplemented with 100 U.mL^−1^ penicillin and 100 µg.mL^−1^ streptomycin (Gibco). Cells were grown at 37°C, 20% O_2_, 5% CO_2_. DT40 and HCT116 cells and derivative clones were routinely confirmed to be negative for mycoplasma contamination.

### Tet-ON inducible system

A Tet-ON system based on the T-REx System (Invitrogen) was used to induce the transcription of *DMD* upon addition of the tetracycline antibiotic. Briefly, the tetracycline-inducible promoter (Tet-promoter) used consists of a CMV promoter into which 2 copies of the Tet-operator 2 (TetO) sequence have been inserted in tandem. In the absence of tetracycline, transcription is repressed by the high affinity binding of the Tet-repressor (TetR) to the TetO. Addition of tetracycline results in the release of TetO by the TetR and derepression of the promoter. The Tet-promoter was amplified from a pcDNA4/TO vector (Invitrogen). The TetR was expressed from a pcDNA6/TR vector (Invitrogen).

### Constructs

Targeting constructs for homologous recombination in DT40 cells were created from LoxP vectors^1^ using standard molecular biology and cloning protocols. ≈2 kb-long 5’ and 3’ homology regions were amplified from DT40 genomic DNA with primers listed in Supplementary Table 2. The chicken β-actin promoter was synthesized by GeneScript. All constructs were checked for mutations by sequencing. Guide RNAs targeting *FHIT* promoter (gRNA1: GCCAAATGCCATGTGGGTGC; gRNA2: TCAATTTAGATTTCGGCTTC) were designed using the gRNA design tool from DNA2.0. gRNA/Cas9 plasmid containing sequences for the two gRNAs and wtCas9 was synthesized by DNA2.0. ≈1 kb-long 5’ and 3’ homology regions flanking the CRISPR/Cas9 cutting sites were amplified from RP11-137N22 BAC with primers listed in Supplementary Table 2 and cloned into the HR710PA-1 plasmid (System Biosciences) on each side of the EF1α promoter-hygromycin selection cassette. Detailed cloning procedures are available upon request.

### Transfection and genotyping

10^7^ exponentially growing DT40 cells were electroporated with 35 μg of the linearized construct using a Biorad electroporator set at 25 μF and 550 V for targeted integrations, 960 μF and 250 V for the random insertion of the pcDNA6/TR vector. Recombinant clones were selected with either 21.75 μg.mL^−1^ blasticidin, 0.5 μg.mL^−1^ puromycin or 450 μg.mL^−1^ zeocin, identified by PCR using LA Taq DNA Polymerase (Takara) and confirmed by Southern blot. At least two positive clones were randomly selected and amplified for further studies. Primers used for genotyping are listed in Supplementary Table 2. Probes used for Southern blot were amplified from the targeting constructs using primers listed in Supplementary Table 2. The 2-log DNA ladder (NEB) was used as molecular-weight size marker for agarose gel electrophoresis except in Supplementary Fig. 1f where the GeneRuler 1 kb Plus DNA ladder (ThermoFisher Scientific) was used.

For transient transfection, 3×10^6^ exponentially growing DT40 cells were transfected with 5 μg of a reporter plasmid expressing the luciferase gene under the control of the Tet-promoter using the Amaxa Nucleofector system (T solution, program B-023).

HCT116 cells were co-transfected with 1 µg gRNA/Cas9 plasmid and 1µg HR710PA-1-*FHIT* plasmid using FuGENE HD Transfection Reagent (Promega) according to the manufacturer’s instructions. Recombinant cells were selected with 200 µg.mL^−1^ hygromycin B (Invitrogen) 48 h post-transfection. 2 µM ganciclovir was added to the culture medium 7 days post-transfection to counter-select cells containing randomly integrated HR710PA-1-*FHIT* plasmid. After a further 12 days, single clones were selected and amplified. Positive clones were identified by PCR and confirmed by sequencing.

### Luciferase reporter assay

DT40 clonal cell lines expressing the Tet-repressor were tested for their ability to induce transcription of the luciferase reporter gene upon addition of tetracycline, which was quantified by the light emitted during an enzymatic reaction of bioluminescence using the Dual-Glo Luciferase Assay System (Promega) according to the manufacturer’s instructions. Luminescence was measured using a luminometer 24 h after addition of 1 μg.mL^−1^ tetracycline (SIGMA) to the culture medium.

### Tetracycline-induced transcriptional activation of *DMD*

Transcriptional activation of *DMD* was achieved with a 24 h treatment of DMD^Tet/Tet^ and DMD^+/Tet^ cells with 1 μg.mL^−1^ tetracycline. A low transcriptional background from the Tet-promoter is detected even without tetracycline (Figs 1a and 3a), as already observed^2^.

### Floxed cassette excision

The DT40 cell line used in this study contains a stably integrated MerCreMer plasmid^1^. The protein is inactive in the absence of 4-hydroxytamoxifen (4-OHT) due to the hormone binding domains of the Mutated estrogen receptor (Mer) fused either side of the Cre recombinase. For excision of floxed cassettes, cells were cultured for 24 h with 0.5 μM 4-OHT then distributed into 96-well flat-bottom microtiter plates at a concentration of 1, 3 and 30 viable cells per well. Isolated colonies were tested for cassette excision by loss of antibiotic resistance and PCR.

### mRNA quantification

DT40 cells were collected during their exponential growth phase. Total RNA was extracted using miRNeasy Mini Kit (Qiagen) according to the manufacturer’s instructions and treated with DNase (Roche). 1 μg of total RNA was reverse-transcribed using the iScript cDNA Synthesis Kit (Biorad). Real-time quantitative PCR reactions were set up using GoTaq qPCR Master Mix (Promega) and run on a LightCycler 480 II (Roche). Each reaction was performed in triplicate. DNA contamination was quantified in reverse transcriptase free reactions. Primer sequences are listed in Supplementary Table 3. Each primer pair spanned one intron to avoid unwanted amplification of genomic DNA. The efficiency of each primer pair was tested by performing qPCR with the same protocol on increasing dilutions of cDNA (from 1/10 to 1/10,000 dilutions) and calculated using the coefficient of amplification. The efficiency of each pair being similar, it is possible to directly compare the amount of each amplification product. mRNA levels were quantified relative to β-actin (*ACTB*) mRNA. For DMD^Tet/Tet^, DMD^+/Tet^ and DMD^+/βa^ cells, mRNA levels were quantified before the excision of the floxed selection cassettes. For each construct, mRNA quantification experiments were performed using at least two different clonal cell lines.

Total RNA from HCT116 cells was isolated using RNeasy Mini Kit (Qiagen) according to the manufacturer’s instructions. cDNA was synthesized with Maxima First Strand cDNA Synthesis Kit for RT-qPCR (ThermoFisher Scientific). Real-time quantitative PCR was performed using Luminaris Color HiGreen qPCR Master Mix, low ROX (ThermoFisher Scientific) on a 7500 Real-Time PCR System (Applied Biosystems). Primer sequences are listed in Supplementary Table 3. *FHIT* mRNA levels were quantified relative to RNA polymerase II subunit F (*POLR2F*), ribosomal protein L11 (*RPL11*) and cyclophylin B (*PPIB*) mRNA. mRNA quantification experiments were performed using one FHIT^EF1α/EF1α^ clonal cell line.

### Quantification of 5-Ethynyl uridine (EU) incorporation into nascent RNAs

10^7^ exponentially growing DT40 cells were treated for 1 h with 0.5 mM EU. EU-labelled RNA was captured using the Click-iT Nascent RNA Capture kit (Invitrogen) according to the manufacturer’s instructions. Briefly, 1 μg of total RNA was biotinylated thanks to the click reaction between EU and azide-modified biotin, then purified using streptavidin magnetic beads. Nascent RNAs were reverse-transcribed on the beads using Superscript kit (Invitrogen). Real-time quantitative PCR reactions were performed as described above for DT40 samples using intra-intronic primers listed in Supplementary Table 3. All primers had similar efficiencies. RNA levels were calculated relative to β-actin mRNA quantities.

### Fluorescent *in situ* hybridization (FISH) on metaphase chromosomes

DT40 and HCT116 cells were grown for 16 h with 0.6 and 0.15 μM aphidicolin, respectively. Metaphase spreads were prepared according to standard cytogenetic procedures after a 3 h treatment with 0.1 μg.mL^−1^ colcemid for DT40 cells and a 2 h treatment with 100 nM nocodazol for HCT116 cells. For untreated DT40 cells, metaphase spreads were prepared after a 1.5 h treatment with 0.1 μg.mL^−1^ colcemid.

For DT40 FISH analyses, probes correspond to adjacent 5 kb-long PCR products spread over ≈50 kb delimiting *DMD*, *CCSER1* or *PARK2* genes (see probe coordinates in Supplementary Table 4); probes were amplified from DT40 genomic DNA with primers listed in Supplementary Table 4. Probes were labelled either with biotin using the BioPrime DNA labelling system (Invitrogen) or with digoxigenin using the DIG DNA labelling mix (Roche) and subsequently purified on Illustra ProbeQuant G-50 Micro Columns (GE Healthcare).

FISH was performed essentially as described^3^ without the proteinase K step and using 75 ng of each probe. Chicken Hybloc DNA (Amplitech) was used as a repetitive sequence competitor DNA. Immunodetection was performed by successive incubations in the following reagents: (i) for biotinylated probes, (1) Alexa Fluor 555-conjugated streptavidin (Invitrogen), (2) biotin-conjugated rabbit anti-streptavidin (Rockland Immunochemicals), (3) Alexa Fluor 555-conjugated streptavidin (Invitrogen) (ii) for digoxigenin-labelled probes, (1) FITC-conjugated mouse anti-digoxin (Jackson ImmunoResearch), (2) Alexa Fluor 488-conjugated goat anti-mouse (Invitrogen). Chromosomes were counterstained with 49,6-diamidino-2-phenylindole (DAPI) (Vectashield mounting medium for fluorescence with DAPI; Vector Laboratories) and metaphases were observed by fluorescence microscopy. Aphidicolin-induced breaks at the 679.5 kb-long *PARK2* gene were used as a control to demonstrate that modulation of *DMD* or *CCSER1* transcription specifically impacts the fragility of those genes and that the limited perturbations of cell physiology observed in CCSER1^βa/βa^ cells (Supplementary Fig. 4d, e) do not perturb break induction by aphidicolin. For each construct, FISH experiments were performed using at least two different clonal cell lines.

FISH analysis of FRA3B fragility in HCT116 cells was performed using 50 ng of biotinylated RP11-641C1, RP11-32J15 and RP11-147N17 BACs selected from the human genome project RP11 library. Immunodetection was performed by successive incubations in Alexa Fluor 488-conjugated streptavidin (Invitrogen) and biotin-conjugated rabbit anti-streptavidin (Rockland Immunochemicals). 15 µg Cot-1 DNA (Invitrogen) were used as a repetitive sequence competitor DNA. FISH experiments were performed using one FHIT^EF1α/EF1α^ clonal cell line.

### Bivariate Fluorescence-Activated Cell Sorting (FACS) analysis

Cell cycle analyses were performed as follows. Exponentially growing DT40 cells were pulse-labelled for 15 min with 30 μM bromodeoxyuridine (BrdU) and fixed in ethanol. After partial denaturation of DNA following HCl/pepsin treatment, immunodetection was performed by incubations with rat anti-BrdU (Bio-Rad, formerly AbD Serotec) then Alexa Fluor 488-conjugated chicken anti-rat (Invitrogen). DNA was counterstained with propidium iodide in the presence of 25 μg.mL^−1^ RNase. Samples were analyzed using a BD Biosciences LSRII flow cytometer with BD FACSDiva software. Data were processed using FlowJo v8.7.3. Gating strategy is illustrated in Supplementary Fig. 7.

### Cell growth and doubling time

The cumulative growth curves of the DT40 cell lines used in this study were determined after normalization for dilution at each subculture. Doubling times were estimated based on the growth curves. Indeed, for exponentially growing cells, the final number of cells (N) is given by the formula N = initial number of cells x 2^number of doublings^, where the number of doublings corresponds to the duration of culture divided by the doubling time (T); it follows that T = [duration of culture*log(2)]/[log(final number of cells)-log(initial number of cells)].

### Replication timing analyses

Timing analyses in DT40 cells were made as previously described^4^. Briefly, exponentially growing DT40 cells were pulse-labelled for 1 h with 3 mM BrdU, fixed in ethanol and stored at −20°C overnight. Fixed cells were re-suspended in 1X PBS with 50 μg.mL^−1^ propidium iodide and 1 mg.mL^−1^ RNase, and incubated for 30 min at room temperature. Cells were sorted by flow cytometry based on their nuclear content using a BD Biosciences INFLUX cell sorter with the BD FACS™ Sortware software (v 1.0.0.650). S-phase was divided into four fractions from early to late S-phase named S1 to S4. 50,000 cells were sorted in each fraction. DNA was extracted by phenol/chloroform, sonicated to obtain fragments between 500-1000 bp in size, and BrdU-labelled DNA was immunoprecipitated using mouse anti-BrdU antibody (BD Biosciences). Real-time quantitative PCR was performed using the Roche LightCycler 2.0 detection system with the Absolute QPCR-SYBR Green mix (ThermoFisher Scientific). For each reaction, amplification of the purified BrdU-labelled DNA was performed in duplicate. As mitochondrial DNA replicates throughout the cell cycle and should be equally represented in every fraction, the amount of immunoprecipitated DNA in each S-phase fraction was normalized by the abundance of mitochondrial DNA measured using a specific primer pair (MIT). Quality control experiments were performed to confirm enrichment of known early-, mid- and late-replicated loci in the expected fractions (Supplementary Table 5). In heterozygous cells, primer pairs overlapping the site of insertion and next to the site of insertion of the Tet- or β-actin promoters were used to detect the timing of the wild type allele (“without” primers) and both alleles (“both” primers), respectively. A primer pair specific to the transgene (“with” primers) was used to analyze the timing of the modified allele. For each construct, replication timing analyses were performed on two different clonal cell lines.

For replication timing analyses of *FHIT*, exponentially growing HCT116 cells were pulse-labelled for 1 h with 50 μM BrdU, fixed in ethanol and incubated overnight at −20°C in the presence of 15 μg.mL^−1^ Hoescht 33342 (ThermoFisher Scientific). Fixed cells were re-suspended in 1X PBS and cells were sorted by flow cytometry based on their nuclear content using a BD Biosciences INFLUX cell sorter. S-phase was divided into four fractions, from early to late S-phase (S1 to S4). To check the quality of sorted HCT116 fractions, the post-sort cells, already stained with Hoescht 33342, were directly re-analyzed by flow cytometry. DNA was isolated using Maxwell RSC Blood DNA kit (Promega) according to the manufacturer’s instructions, and sonicated. For each fraction, immunoprecipitation of BrdU-labelled DNA was performed on 5 μg of DNA using mouse anti-BrdU antibody (BD Biosciences). Immunoprecipitated DNA was further extracted by phenol/chloroform, and real-time quantitative PCR was performed using QuantiNova SYBR Green PCR kit (Qiagen) on a 7300 Real-Time PCR System (Applied Biosystems). Replication timing analyses were performed using one FHIT^EF1α/EF1α^ clonal cell line.

All primer pairs used for replication timing analyses are listed in Supplementary Table 6.

### DNA-combing

Neo-synthesized DNA was labelled as described^5^ with the following changes: exponentially growing DT40 cells were pulse-labelled for 20 min with 20 μM iododeoxyuridine (IdU) followed by a 20 min pulse with 100 μM chlorodeoxyuridine (CldU), then by a 5 min chase with 1 mM thymidine. Genomic DNA was extracted and combing was performed on silanized coverslips prepared by plasma cleaning and liquid-phase silanization as described^6, 7^ using a Genomic Vision apparatus.

### FISH on combed DNA and immunofluorescence detection of neo-synthesized DNA

Morse-codes were designed as described^8^ to specifically identify DNA molecules spanning the *DMD* or *CCSER1* loci. Morse-code probes correspond to a collection of ≈5 kb-long PCR products spread all over the locus of interest, separated by precise distances and divided into voluntarily different and asymmetrical patterns to correctly orient the DNA fibre. Morse-codes for *DMD* and *CCSER1* detection are made of 32 probes and 27 probes, respectively. PCR products were prepared and labelled with biotin as described above for FISH on metaphases. Primer pairs used are listed in Supplementary Table 7. Hybridization of the probes was carried out as described previously^5^. Immunodetection was performed by successive incubations in the following reagents: (1) Alexa Fluor 488-conjugated streptavidin (Invitrogen), (2) biotin-conjugated rabbit anti-streptavidin (Rockland Immunochemicals), (3) Alexa Fluor 488-conjugated streptavidin (Invitrogen), mouse anti-BrdU (BD Biosciences) and rat anti-BrdU (Bio-Rad, formerly AbD Serotec), (4) biotin-conjugated rabbit anti-streptavidin (Rockland Immunochemicals), Alexa Fluor 350-conjugated goat anti-mouse (Invitrogen) and Alexa Fluor 594-conjugated donkey anti-rat (Invitrogen), (5) Alexa Fluor 488-conjugated streptavidin (Invitrogen), Alexa Fluor 350-conjugated donkey anti-goat (Invitrogen) and mouse anti-single stranded DNA (Millipore), (6) Cy5.5-conjugated goat anti-mouse (Abcam) and (7) Cy5.5-conjugated donkey anti-goat (Abcam). For bulk genome analyses, immunodetection was performed as follows: (1) FITC-conjugated mouse anti-BrdU (BD Biosciences) and rat anti-BrdU (Bio-Rad, formerly AbD Serotec), (2) Alexa Fluor 488-conjugated goat anti-mouse (Invitrogen) and Alexa Fluor 555-conjugated goat anti-rat (Invitrogen), (3) mouse anti-single stranded DNA (Millipore), (4) Cy5.5-conjugated goat anti-mouse (Abcam) and (5) Cy5.5-conjugated donkey anti-goat (Abcam). Antibody incubations, washes and slide mounting were performed as reported previously^5^.

### Image acquisition

Images were acquired on a motorized XY stage of an Axio Imager Z2 (Carl Zeiss) or a DM6000 B (Leica) epifluorescence microscope connected to a CoolSNAP HQ2 charge-coupled device camera (Roper Scientific) and run by Metamorph software (Molecular Devices). A X100 objective was used for imaging methaphase chromosomes and a X63 objective was used for imaging combed DNA fibres. For images of the *DMD* and *CCSER1* loci, two overlays of images were set up for each microscope field. The first one combined the IdU/CldU and FISH signals to identify the fibres of interest, replicating or not. The second overlay combined the IdU, FISH and DNA signals to determine the length of the DNA fibre bearing the Morse-code. DNA counterstaining was systematically used to ensure that several fibres are not overlapping (ii) that replication signals belong to the same fibre and (iii) that replication signals are intact.

### Signal treatment and statistical analyses

Measurement of DNA fibre and replication tract length was performed on imaged DNA molecules using Adobe Photoshop CS5.1. DNA fibres bearing Morse-code signals were modelled and aligned on a schematic representation of the *DMD* or *CCSER1* loci using Adobe Illustrator CS5.1. Graphical representations of *DMD* and *CCSER1* were drawn based on the locus length, the theoretical ≈2 kb/μm stretching factor of the combing apparatus indicated by the manufacturer, and the resolution of the camera at a magnification of X63 (0.1024 μm/pixel). For instance, the 996,168 bp *DMD* gene corresponded to 996.168/2/0.1024=4864.1 pixels. Morse-code probes were used to calculate the actual stretching factor of each DNA molecule by comparing the length of the fibre imaged with the microscope with the theoretical length defined according to the position of the probes. Overall, we found a mean stretching factor of 1.93 kb/μm, which is extremely close to the expected value of 2 kb/μm. Still, the stretching factor of individual fibres fluctuated from 1.7 to 2.4 kb/μm. Therefore, to prevent that variations in the stretching of the DNA molecules during the combing step introduce a bias in tract measurement, the lengths of IdU and CldU tracts were normalized by the stretching factor of the DNA fibre on which they were located. For bulk genome analyses, the lengths of IdU and CldU tracts were normalized by the mean stretching factor calculated when analyzing *DMD* or *CCSER1* loci using the exact same sample and coverslip batch. Replication fork speed was then estimated on individual forks as (i) the ratio (l_Idu_+l_CldU_)/(t_IdU_+t_CldU_) for forks displaying an intact IdU tract flanked on one side by an intact CldU tract, (ii) the ratio l_Idu_/t_IdU_ for forks with an intact IdU signal flanked on one side by a broken CldU signal and (iii) the ratio l_Cldu_/t_CldU_ for forks with an intact CldU signal flanked on one side by a broken IdU signal, with l_IdU_ and t_IdU_ being the measured length (in kb) and labelling time (in min) for IdU incorporation, respectively, and l_CldU_ and t_CldU_ the corresponding parameters for CldU incorporation. For all experiments, t_IdU_=t_CldU_=20 min. Replication signal integrity was ascertained by DNA counterstaining. For fork progression analyses in *DMD* and *CCSER1*, only forks overlapping at least partially these loci were taken into account. Coordinates and length of DNA molecules and replication signals were compiled using Microsoft Excel for Mac 2011. Statistical comparisons of fork speed distributions were assessed with the nonparametric Mann-Whitney-Wilcoxon test (two-tailed) using GraphPad Prism 6 (GraphPad Software). No assumptions or corrections were made. Statistical significance was set to p ≤ 0.05. All DNA fibres for CCSER1^βa^ allele originate from one biological sample, DNA combing results for WT *CCSER1* and *DMD* loci compile DNA fibres from two distinct biological WT samples and DNA combing results for DMD^Tet/Tet^ cells compile DNA fibres from two clonal cell lines. Replication analysis of WT and modified *CCSER1* and *DMD* loci by DNA combing was performed once.

### Coverage profile

Graphical representations of “Total DNA” and “DNA with replication” coverages of *DMD* and *CCSER1* were made with R^9^ using custom scripts adapted from^10^.

### Genomic coordinates and genome annotations

Coordinates are given according to the ICGSC/galGal4 chicken genome or the GRCh37/hg19 human genome assemblies. *DMD* is the largest annotated gene of the chicken genome both in the latest RefSeq and Ensembl gene annotations available when this manuscript was written.

### Code availabilty

Custom R scripts are available upon request.

### Data availabilty

The data that support the findings of this study are available upon request.

