## Supplementary Materials for "Transcription-dependent regulation of replication dynamics modulates genome stability"

### Supplementary Figure 1

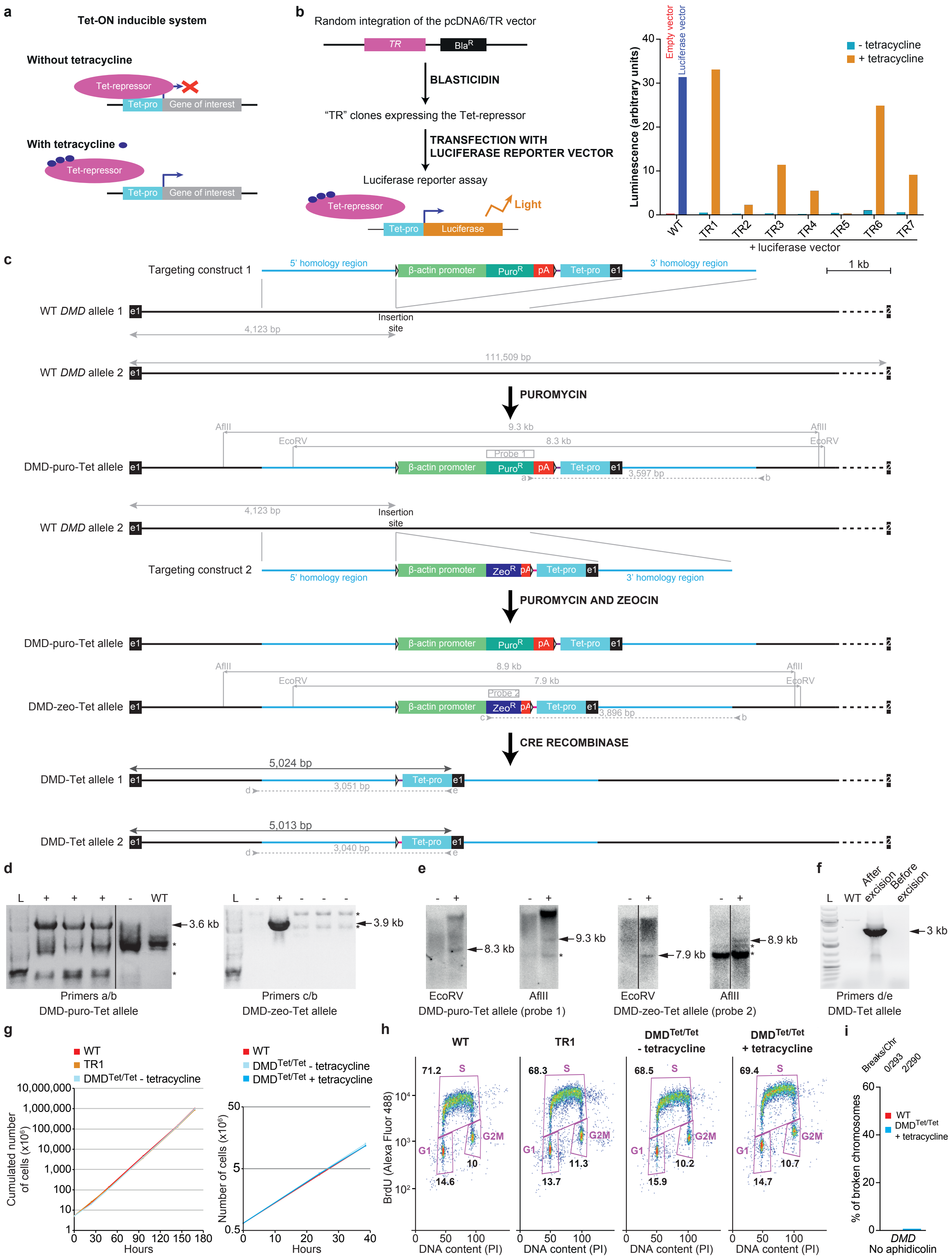

### Supplementary Figure 2

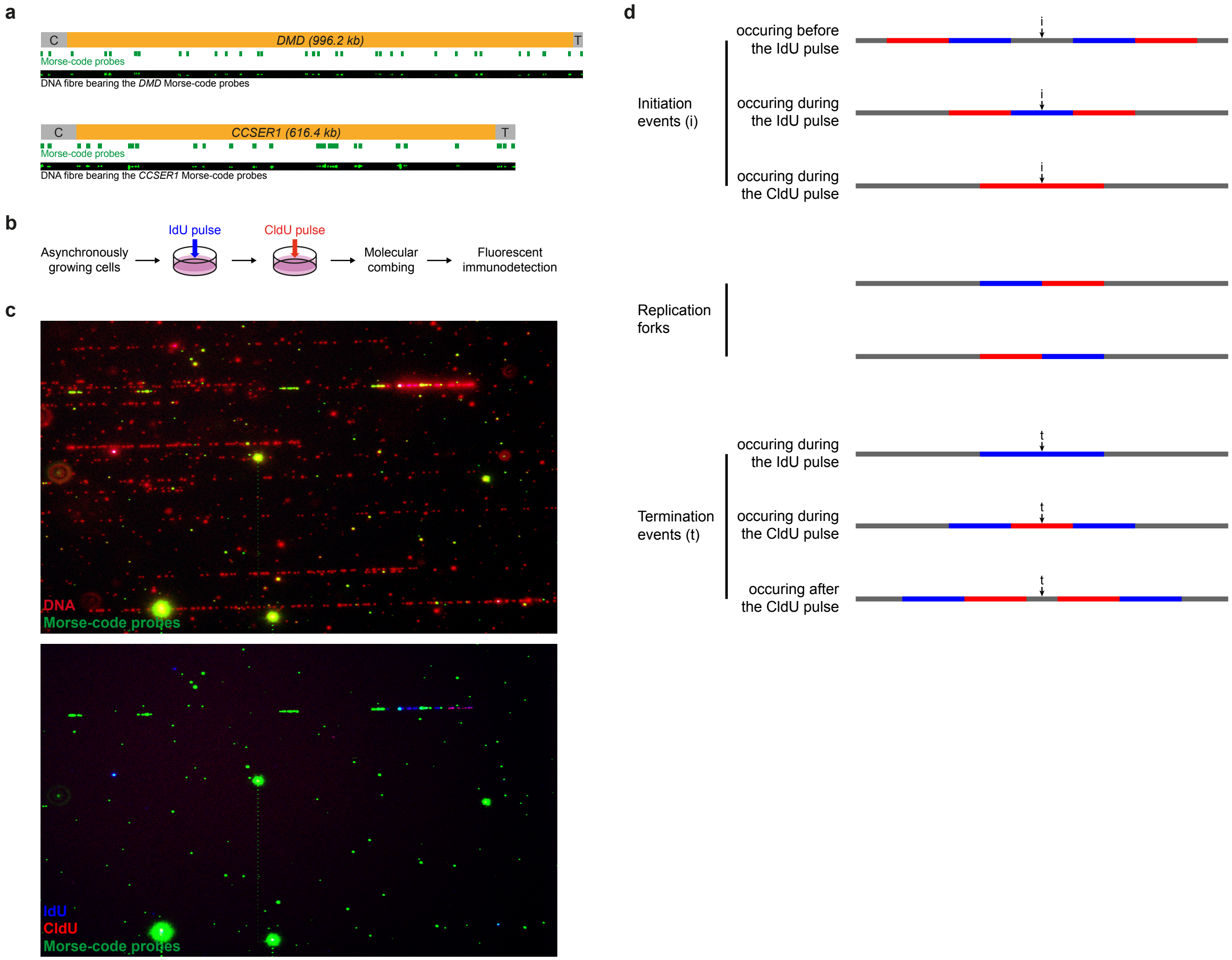

#### Supplementary Figure 3

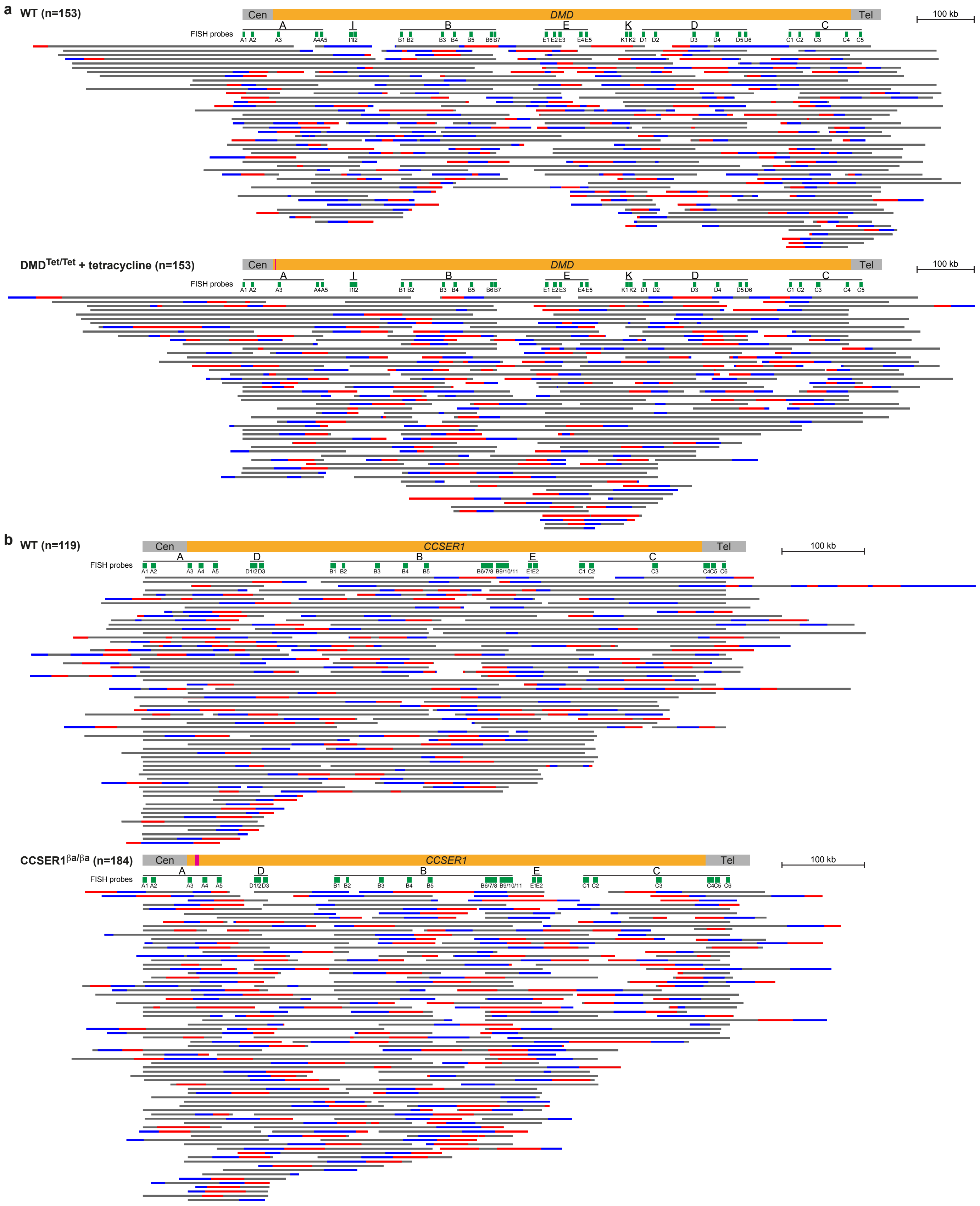

### Supplementary Figure 4

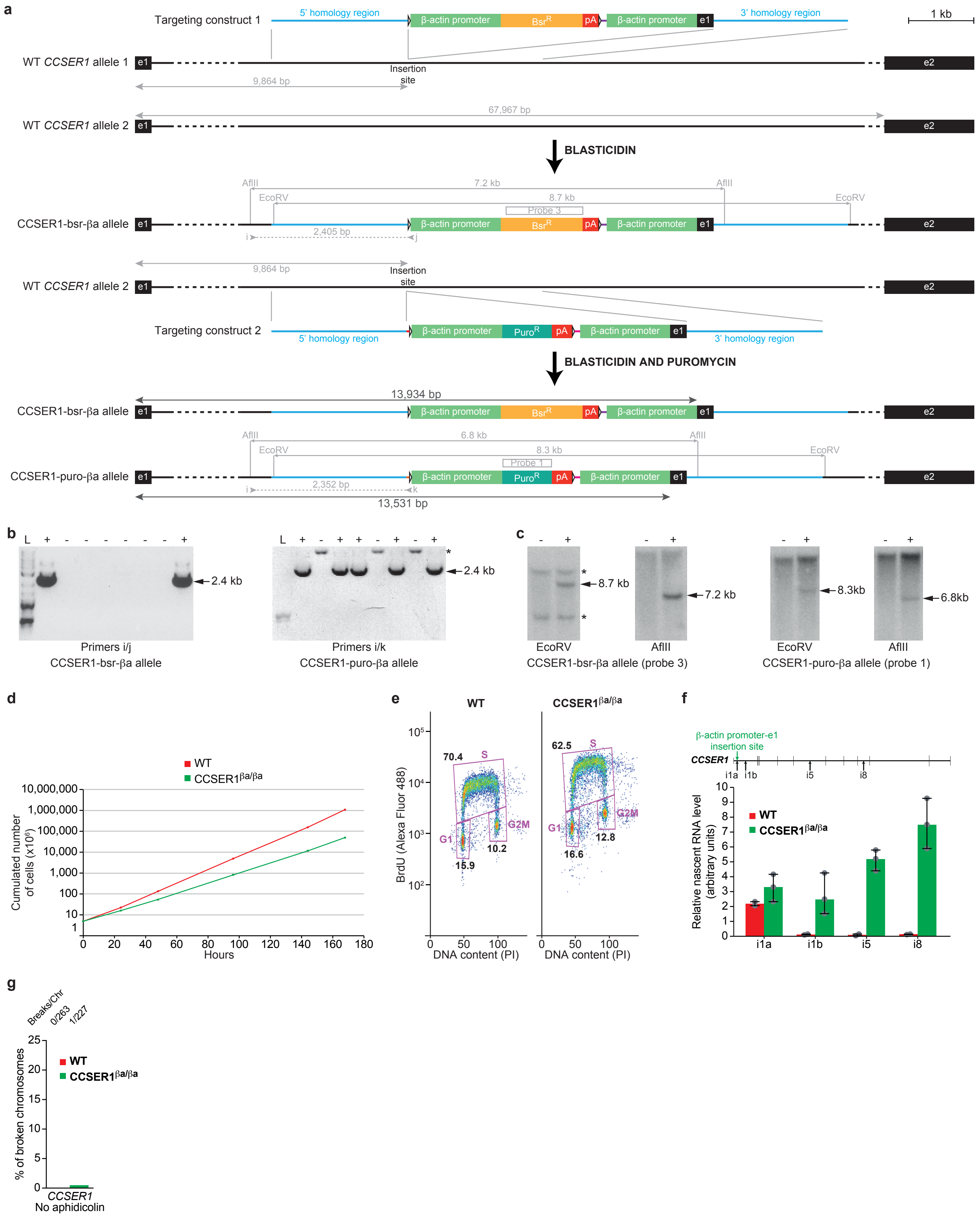

#### Supplementary Figure 5

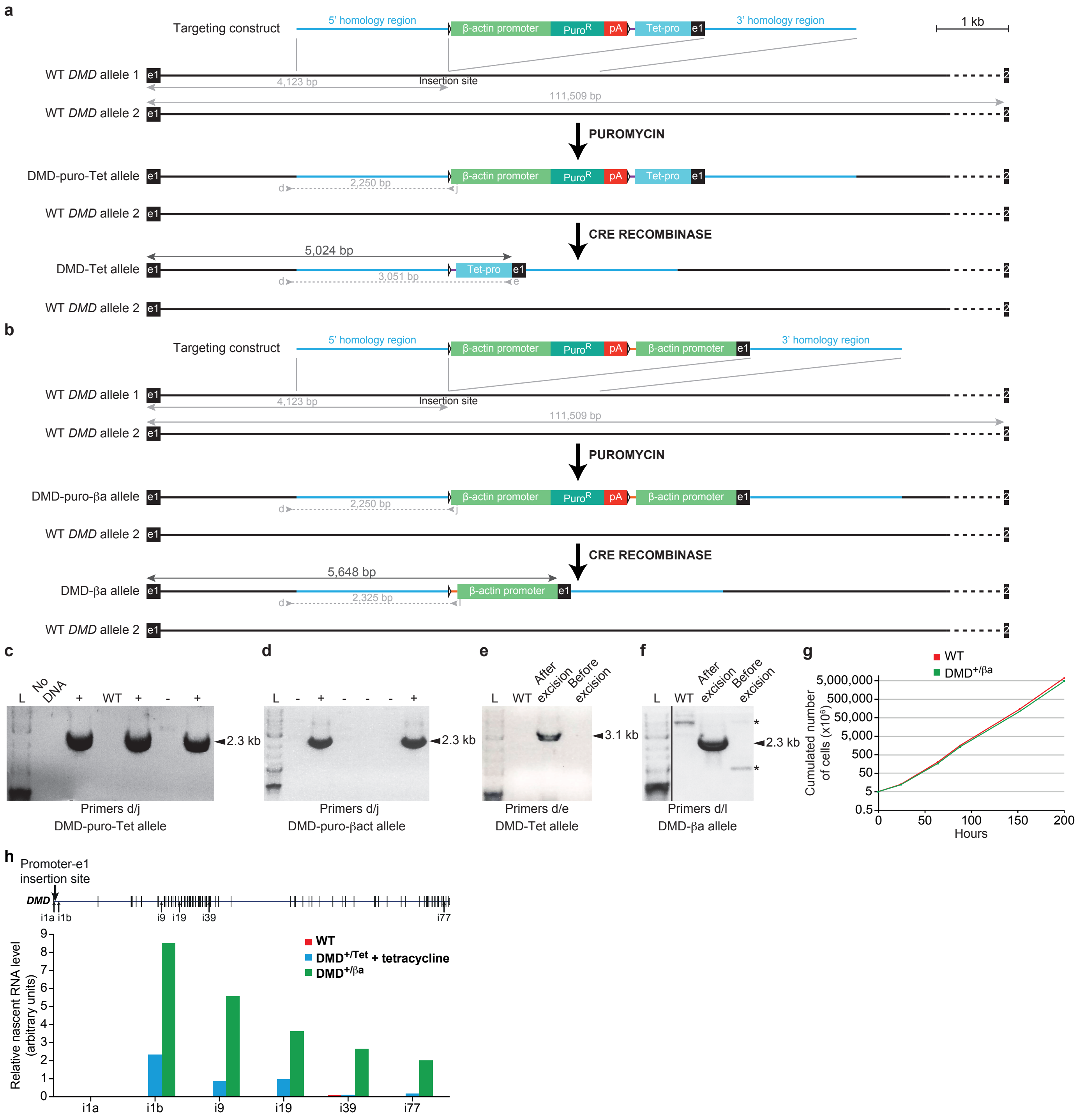

### Supplementary Figure 6

**a**

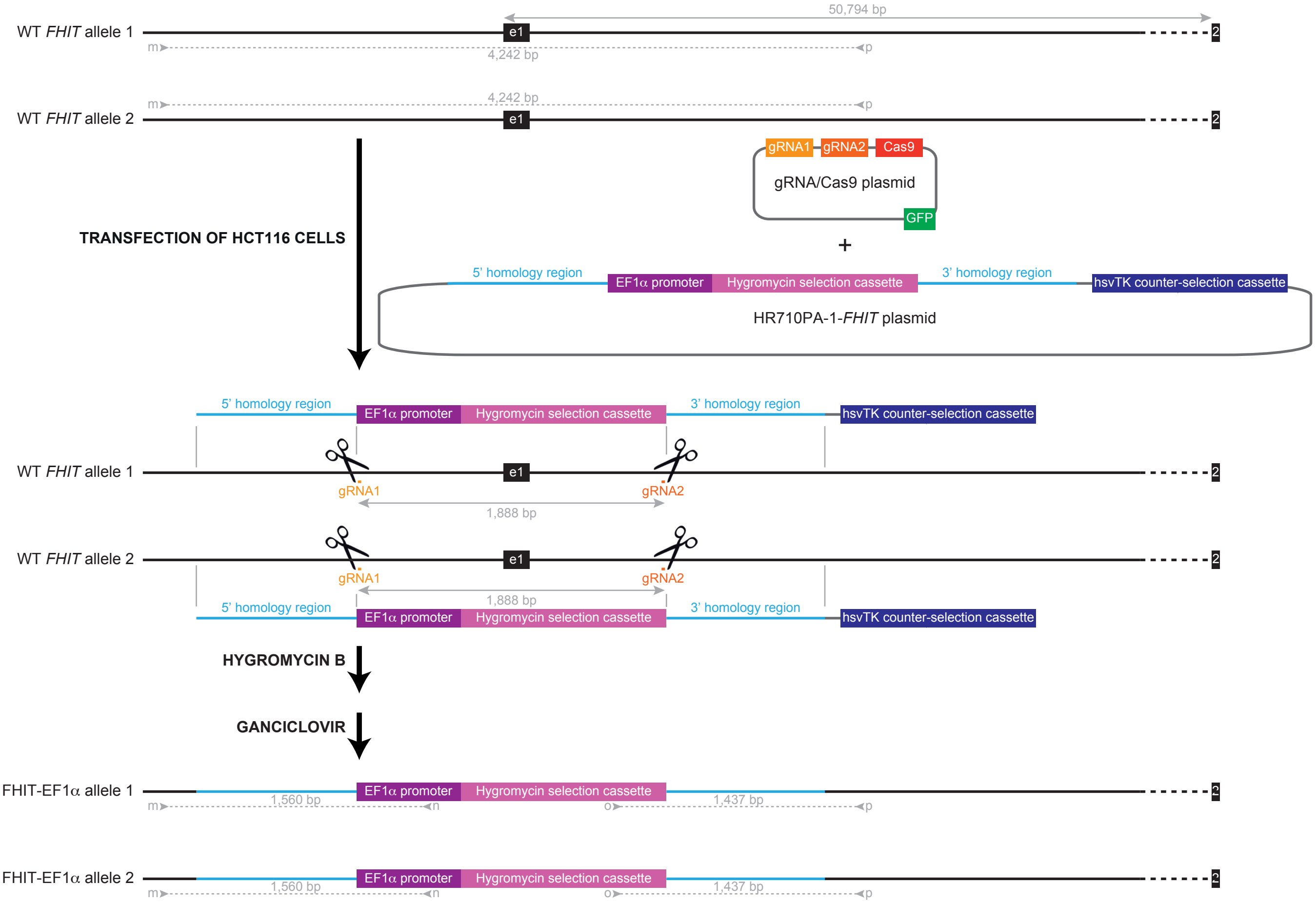


**b**

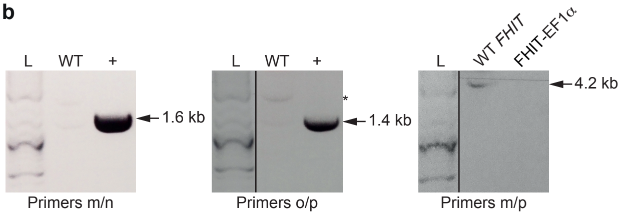

**Supplementary Figure 7**

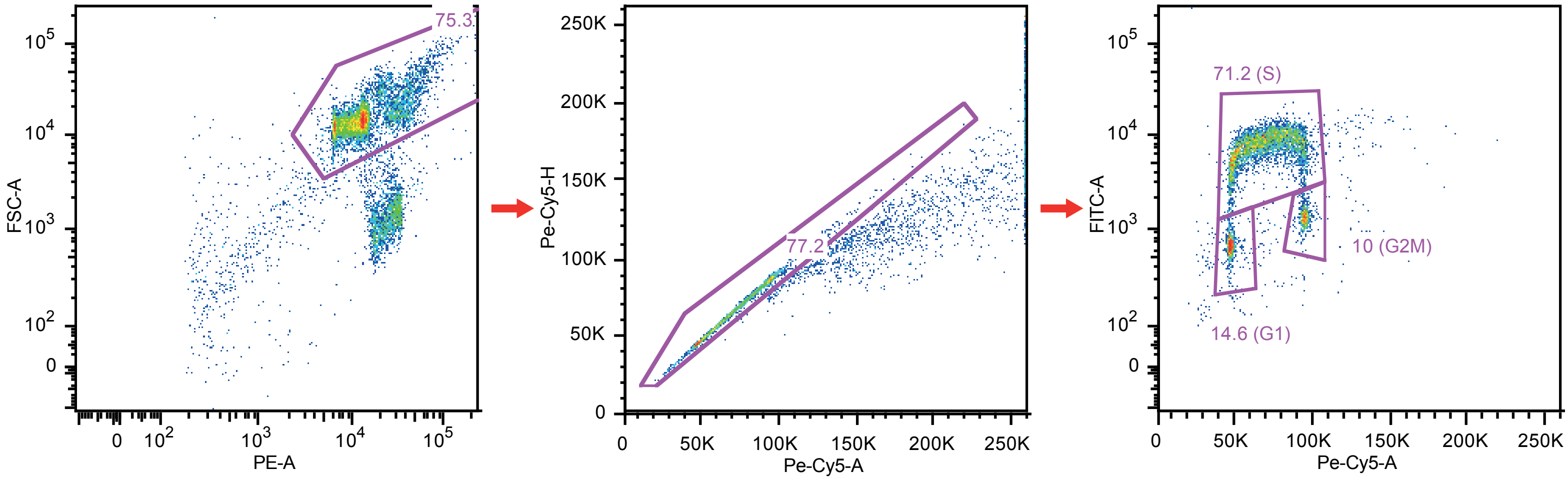


Ungated
WT
Event Count: 10000

Live
WT
Event Count: 7526

Single
WT
Event Count: 5811

#### Supplementary Figure legends

**Supplementary Figure 1. Tetracycline-induced transcriptional activation of *DMD*.**

**a,** Transcriptional activation using a Tet-ON system. Tet-pro, Tet-promoter.

**b,** Construction of a stable DT40 cell line expressing the TR gene encoding the Tet-repressor. Clone TR1 exhibited the best tetracycline-induced transcriptional activation of the luciferase reporter gene and was subsequently used for the Tet-promoter integration. $Bla^R$, blasticidin resistance gene.

**c,** Targeting strategy. The 5' part of the two *DMD* alleles, starting at exon 1 (e1, black box), is shown. Alleles were sequentially targeted in the TR1 cell line with constructs 1 then 2. Despite numerous attempts, we were unable to integrate the Tet-promoter precisely upstream of the first exon of *DMD*; we therefore shifted the insertion site and managed to integrate the construct 4.1 kb downstream of *DMD* transcription start site (TSS). A synthetic *DMD* exon 1 was inserted together with the Tet-promoter to fully reconstitute the *DMD* transcript, which was confirmed by RT-qPCR (see Fig. 1a). As a result, the DMD-Tet allele is ≈5 kb shorter than the WT *DMD* gene (991 and 996 kb, respectively).

Targeting construct 1 contains (1) a 2.1 kb-long 5' homology region, (2) a puromycin selection cassette made of the chicken β-actin promoter, the puromycin resistance gene ($Puro^R$) and SV40 polyadenylation signals (pA) flanked by LoxP sites (white arrowheads), (3) a 71 bp linker (violet line), (4) the Tet-promoter (Tet-pro) upstream of a synthetic *DMD* exon 1, (5) a 2.1 kb-long 3' homology region. Targeting construct 2 is similar to targeting construct 1 except that the selection cassette contains a zeocin resistance gene ($Zeo^R$) and that a 60 bp linker (pink line) was used. Recombinants were identified by PCR using the indicated primers (grey arrows) and by Southern blot with AflII and EcoRV restriction sites and the indicated probes. Excision of the selection cassettes was achieved by transient induction of Cre recombinase activity and verified by PCR with primers d/e.

**d, e,** Screening of recombinant clones by PCR (**d**) and Southern blot (**e**). + and - indicate positive and negative clones, respectively. Arrows point to the bands of the expected size described in **c**. Asterisks (*) indicate non-specific bands. L, ladder.

**f,** Verification of the excision of the floxed selection cassettes by PCR using primers d/e. Before excision, the presence of the GC-rich (>71.76%) β-actin promoter prevents amplification in standard PCR conditions. L, ladder.

**g, h,** Activation of *DMD* transcription has no visible impact on cell growth nor cell cycle.

**g,** Representative growth curves of WT, TR1 and $DMD^{Tet/Tet}$ cells without tetracycline (left panel) and of WT and $DMD^{Tet/Tet}$ cells with or without tetracycline (right panel). The population doubling time of both WT cells and $DMD^{Tet/Tet}$ cells with tetracycline is ≈9.2 h, indicating that the induction of *DMD* transcription has no impact on cell growth.

**h,** Representative bivariate BrdU/DNA FACS analysis of WT, TR1 and $DMD^{Tet/Tet}$ cells with or without tetracycline. Gates indicate the fraction of cells in the G1, S and G2+M phases of the cell cycle.

**i,** *DMD* is not fragile in the absence of aphidicolin in WT and $DMD^{Tet/Tet}$ cells with tetracycline. Metaphase spreads were prepared from exponentially growing cells. Breaks were localized by FISH using probes flanking *DMD*. Raw numbers are presented on top.

**Supplementary Figure 2. Molecular combing analysis of the *DMD* and *CCSER1* loci.**

**a,** Upper panel: *DMD* gene (yellow box) and Morse-code probes (green boxes) used in DNA-combing experiments to identify the *DMD* locus. An example of a DNA fibre bearing the whole Morse-code is shown (the DNA fibre presented here spans several microscope fields of view; please note that background was removed to improve clarity). Lower panel: same as above for the *CCSER1* gene. C, centromeric; T, telomeric.

**b,** Scheme of the protocol for double pulse-labelling of replication in asynchronous cell cultures.

**c,** Example of raw images showing combed DNA fibres from WT DT40 cells. Upper image: DNA fibres counterstained in red and Morse-code probes in green. Please note that the red channel was adjusted. Lower image: same microscope field with IdU and CldU replication signals revealed in blue and red, respectively, and Morse-code probes in green. A fibre bearing part of the *DMD* Morse-code and a replication signal consisting of an IdU tract followed by an interrupted CldU tract is detected.

**d,** Principles of replication signal analysis. A schematic representation of typical replication patterns visualized after immunofluorescence is shown. Blue and red tracts represent neo-synthesized DNA labelled with IdU and CldU, respectively. DNA fibres are in grey. Successive pulses of IdU and CldU allow the recognition of ongoing replication forks, initiation and termination events. Black vertical arrows indicate the estimated positions of initiation and termination events.

**Supplementary Figure 3. Schematic representation of all DNA fibres with replication signals analyzed in this study.**

**a, b,** From top to bottom: *DMD* (**a**) or *CCSER1* (**b**) loci (yellow box), Morse-code probes (green boxes) used in DNA-combing experiments and DNA fibres aligned along the locus using the Morse-code. Morse-codes comprise 32 probes organized in seven motifs (A-E, I, K) and 27 probes organized in five motifs (A-E) for *DMD* and *CCSER1*, respectively. Pink boxes represent the inserted constructs. DNA fibres are represented by a grey line, IdU and CldU tracts are in blue and red, respectively. Only DNA fibres with replication signals are shown. The cell line and number of fibres with replication signals (n) are indicated on the left. DNA fibres are randomly organized. Please note that DNA fibres are fairly often interrupted at the level of a Morse-code probe; this is especially visible for fibres aligned along the *CCSER1* locus. The explanation stems from the frequently dotted DNA fibre counterstaining that prevents a precise determination of the DNA molecule boundaries, which were therefore set to the visible Morse-code probes. This also explains why the total DNA coverage is not entirely homogeneous bus is higher at the level of the Morse-code probes used to identify *DMD* or *CCSER1* (see Figs 1c, d and 2c, d). Cen, centromeric; tel, telomeric.

**Supplementary Figure 4. Upregulation of *CCSER1* transcription.**

**a,** Targeting strategy. The 5' part of the two *CCSER1* alleles, starting at exon 1 (e1, black box), is shown. Alleles were sequentially targeted with constructs 1 then 2. Like for *DMD*, despite numerous attempts, we were unable to replace the endogenous *CCSER1* promoter by the chicken β-actin promoter; we therefore shifted the insertion site and managed to integrate the construct 9,864 bp downstream of *CCSER1* TSS. A synthetic *CSSER1* exon 1 was inserted together with the β-actin promoter to fully reconstitute the *CCSER1* transcript, which was confirmed by RT-qPCR (see Fig. 2a). Moreover, the selection cassette located upstream of the β-actin promoter-exon 1 was not excised in order to block transcription coming from the endogenous *CCSER1* promoter. As a result, the CCSER1-βa allele is ≈14 kb shorter than the WT *CCSER1* gene (602 and 616 kb, respectively). Targeting construct 1 contains (1) a 2 kb-long 5' homology region, (2) a blasticidin selection cassette made of the chicken β-actin promoter, the blasticidin resistance gene ($Bsr^R$) and SV40 polyadenylation signals (pA) flanked by LoxP sites (white arrowheads), (3) a 89 bp linker (violet line), (4) the chicken β-actin promoter upstream of a synthetic *CCSER1* exon 1, (5) a 2 kb-long 3' homology region. Targeting construct 2 is similar to targeting construct 1 except that the selection cassette contains a puromycin resistance gene ($Puro^R$), a 86 bp linker (pink line) was used, and that a 20 bp sequence (orange line) was inserted between the 5' homology region and the left LoxP site to allow the hybridization of primer k in order to discriminate CCSER1-puro-βa and CCSER1-bsr-βa alleles. Recombinants were identified by PCR using the indicated primers (grey arrows) and by Southern blot with AflII and EcoRV restriction sites and the indicated probes.

**b, c,** Screening of recombinant clones by PCR (**b**) and Southern blot (**c**). + and - indicate positive and negative clones, respectively. Arrows point to the bands of the expected size described in **a**. Asterisks (*) indicate non-specific bands. L, ladder.

**d, e,** Impact of *CCSER1* overexpression on cell physiology.

**d,** Representative growth curves of WT and CCSER1$^{\beta a/\beta a}$ cells. *CCSER1* overexpression slows down cell growth, with a population doubling time of ≈12.4 h compared to ≈9.2 h for WT cells.

**e,** Representative bivariate BrdU/DNA FACS analysis of WT and CCSER1$^{\beta a/\beta a}$ cells. Gates indicate the fraction of cells in the G1, S and G2+M phases of the cell cycle. The S-phase fraction of CCSER1$^{\beta a/\beta a}$ cells is slightly lower than for WT cells.

**f,** Quantification of 5-Ethynyl uridine (EU) incorporation into nascent *CCSER1* RNAs in WT and CCSER1$^{\beta a/\beta a}$ cells (median, extreme values and individual data points). On top, a map of *CCSER1* with its exons, the position of the β-actin promoter-*CCSER1* exon 1 insertion site and that of the intronic primer pairs used to monitor transcript formation is presented. i1a and i1b measure the level of nascent *CCSER1* RNAs upstream and downstream of the promoter insertion site, respectively.

**g,** *CCSER1* is not fragile in the absence of aphidicolin in WT and CCSER1$^{\beta a/\beta a}$ cells. Metaphase spreads were prepared from exponentially growing cells. Breaks were localized by FISH using probes flanking *CCSER1*. Raw numbers are presented on top.

**Supplementary Figure 5. Modulation of *DMD* transcription.**

**a, b,** Targeting strategy for Tet-promoter (**a**) or chicken β-actin promoter (**b**) insertion on one *DMD* allele. The 5' part of the two *DMD* alleles, starting at exon1 (e1, black box), is shown. As previously described (see Supplementary Fig. 1c), constructs were integrated 4.1 kb downstream of *DMD* TSS; a synthetic *DMD* exon 1 was inserted together with the promoter to fully reconstitute the *DMD* transcript, which was confirmed by RT-qPCR (see Fig. 3a). As a result, DMD-Tet and DMD-βa alleles are ≈5 kb shorter than the WT *DMD* gene. **a**, One allele was targeted in the TR1 cell line with the targeting construct 1 described in Supplementary Fig. 1c. **b**, One allele was targeted in WT cells with a targeting construct that contains (1) a 2.1 kb-long 5' homology region, (2) a puromycin selection cassette made of the chicken β-actin promoter, the puromycin resistance gene ($Puro^R$) and SV40 polyadenylation signals (pA) flanked by LoxP sites (white arrowheads), (3) a 89 bp linker (orange line), (4) the chicken β-actin promoter upstream of a synthetic *DMD* exon 1, (5) a 2.1 kb-long 3' homology region. Recombinants were identified by PCR using the indicated primers (grey arrows). Excision of the selection cassettes was achieved by transient induction of Cre recombinase activity and verified by PCR with primers d/e (**a**) or d/l (**b**).

**c, d,** Screening of recombinant clones by PCR in $DMD^{+/Tet}$ (**c**) or $DMD^{+/\beta a}$ (**d**) cells. + and - indicate positive and negative clones, respectively. Arrows point to the bands of the expected size described in **a** and **b**. L, ladder.

**e, f,** Verification of the excision of the floxed selection cassette in $DMD^{+/Tet}$ (**e**) or $DMD^{+/\beta a}$ (**f**) cells by PCR using primers described in **a** and **b**. Before excision, the presence of the GC-rich (>71.76%) β-actin promoter prevents amplification in standard PCR conditions. Asterisks (*) indicate non-specific bands. L, ladder.

**g,** Representative growth curves of WT and $DMD^{+/\beta a}$ cells. Induction of *DMD* transcription by the β-actin promoter has no visible impact on cell growth in heterozygous cells.

**h,** Quantification of EU incorporation into nascent *DMD* RNAs in WT cells, DMD$^{+/Tet}$ cells with tetracycline and DMD$^{+/\beta a}$ cells. Top: Map of *DMD* with its exons and the position of the Tet- or β-actin promoter-*DMD* exon 1 insertion site and of the intronic primer pairs used to monitor transcript formation. i1a and i1b measure the level of nascent *DMD* RNAs upstream and downstream of the promoter insertion site, respectively.

**Supplementary Figure 6. Upregulation of *FHIT* transcription in human HCT116 cells.**

**a,** Targeting strategy. *FHIT* promoter was replaced in HCT116 cells by an EF1α promoter-hygromycin selection cassette using CRISPR/Cas9-mediated genome editing. EF1α (Elongation Factor-1 alpha) is a constitutive, robust promoter of human origin that efficiently can drive ectopic gene expression. The 5' part of the two *FHIT* alleles, starting at exon 1 (e1, black box), is shown. Cas9-mediated cleavage (represented as black scissors) was achieved thanks to a pair of guide RNAs (gRNA1 and gRNA2, light and dark orange lines), resulting in the deletion of a ≈1.9 kb region centered on *FHIT* promoter/exon 1. The inserted EF1α promoter-hygromycin selection cassette was present in an Homologous Recombination (HR) targeting vector (HR710PA-1-*FHIT* plasmid), where it was flanked by sequences homologous to the *FHIT* promoter region 10 bp-adjacent to the Cas9 cutting-sites (the 5' and 3' homology regions extend over 985 and 974 bp, respectively). The HR710PA-1-*FHIT* plasmid also contained a hsvTK selection cassette located next to the 3' homology region, which allowed to eliminate cells that have integrated the plasmid not *via* HR by sensitizing them to ganciclovir. Recombinants were identified by PCR using the indicated primers (grey arrows). The insertion of the EF1α promoter-hygromycin selection cassette massively upregulated *FHIT* expression despite the absence of exon 1, as observed by RT-qPCR (see Fig. 4a).

**b,** Identification of recombinant clones by PCR. + indicates a positive clone. A 4,242 bp-long PCR product is amplified from WT *FHIT* allele using m/p primer pair, while no PCR product is obtained after the insertion of the EF1α promoter-hygromycin cassette, allowing to discriminate between homozygous and heterozygous clones. Arrows point to the bands of the expected size described in **a**. Asterisk (*) indicates a non-specific band. L, ladder.

**Supplementary Figure 7. Gating strategy for bivariate FACS analysis.**

Exponentially growing DT40 cells were pulse-labelled with BrdU and fixed in ethanol. Immunodetection of BrdU was performed after partial denaturation of DNA. DNA was counterstained with propidium iodide (PI) prior to flow cytometry analysis. Cells were gated through the FSC-A versus PE-A (SSC) plot to distinguish between dead and living cells. Living cells were then interrogated by the ratios of area (Pe-Cy5-A) to height (Pe-Cy5-H) of the PI signal to gate out cell aggregates. Finally, living, single cells were analysed for their BrdU uptake (FITC-A) versus their PI signal (Pe-Cy5-A), which provides a typical horseshoe-shaped profile allowing to discriminate between cells in G1, S and G2/M phases of the cell cycle. Pink numbers correspond to the percentage of gated cells.

| Locus | Cells | Number of fibres (with replication) | Mean fibre length (with replication) in kb | Median fibre length (with replication) in kb |
|---|---|---|---|---|
| *DMD* | WT | 1,064 (153) | 276 (271) | 203 (226) |
| | $DMD^{Tet/Tet}$ + tetracycline | 1,018 (153) | 231 (289) | 189 (232) |
| *CCSER1* | WT | 636 (119) | 224 (282) | 178 (242) |
| | $CCSER1^{\beta a/\beta a}$ | 1,436 (184) | 151 (183) | 137 (170) |

**Supplementary Table 1. Parameters of the DNA fibres analyzed in this study.**

| Name | Forward | Reverse |
|---|---|---|
| *DMD* 5 'homology region | CCTTATCTGCACTGTGTTAC | GCAAGGCTTTGTAGTATCTG |
| *DMD* 3' homology region | TCTGTGCTTCTTGCTGTATC | TCCTGACTTCAGCAATACTC |
| *CCSER1* 5' homology region | GAAAGCAGTTACCCTATGAG | GCCTGAACAAATCAATTCTC |
| *CCSER1* 3' homology region | CCGTATCCCAGATTTGTTTG | TCCCACCAATATTTGGTTAC |
| Probe1 | CCTGCAGGTCGGCCGCCA | CCATGGGGTCGTGCGCTCCTTT |
| Probe2 | TAAACCATGGCCAAGTTGAC | TAGAGGTCGACGGTATACAG |
| Probe3 | TAGTAGAAGTAGCGACAGAG | CTCTAGTCAAGGCACTATACA |
| Primer a | CAGCGCCCGACCGAAAGGAGCGCACGACC | |
| Primer b | | TGGCTCAGACTTTGTAAACC |
| Primer c | TAAACCATGGCCAAGTTGAC | |
| Primer d | AGATTTCCTCACCTGTATCC | |
| Primer e | | TTAAACGCTAGAGTCCGGAG |
| Primer i | TCTTGGTCAAATGGGTTTC | |
| Primer j | | AGCCCTGATCAATAACTTC |
| Primer k | | GATACCCAGCTTTCTTGTAC |
| Primer l | | CCGTCGACTACAACTTTATT |
| *FHIT* 5 'homology region | ACCTTGAGCCCAGCAGTCAGCATTATCC | CATTGATTCAGACCCAATTCCTAACC |
| *FHIT* 3' homology region | TGTGAAAGTTTACCACTTGCAGAT | GCCAGAAAGGTTGGACTCCT |
| Primer m | CCTGTCTCCAGATCTGCTTCT | |
| Primer n | | TCTCTAGGCACCCGTTCAAT |
| Primer o | CTCAAAGAGCAGCGAGAAGC | |
| Primer p | | TGGGCCAGTTTCCTAATCTC |

**Supplementary Table 2. Primers used for the creation of targeting constructs and for genotyping.**

| Gene | Name | Forward | Reverse |
|---|---|---|---|
| *DMD* | e1/e2 | GTCTGCTCACGTGCTTTGGTAT | CATTTCGTGAATGTTTTCTTTTGAAC |
| | e19/e20 | TCTCAGATCTCAGAGAAAGAGTCAATG | TGCTGGCATCTTGCAGCTT |
| | e30/e31 | GCAGCAGGAGGCTGTGAGA | AGCGGAGGGTTTTGTCAGTCT |
| | e44/e45 | CCCAATGGGAGAAAGTTAACAAGA | CCACTTTTCCTTGGATTTGTCAA |
| | e74/e75 | TCTGGAGAGTGAAGAAAGAGGTGAA | TCCGCTTGCAAGTTTCGAT |
| | e76/e77 | AGACAGCAGTCAGCCAATGCT | TCGTCCTCGCCCATGGT |
| | i1a | AGAGCACACTCTAGAACCAACTACATG | CAAGTTCTCCTTATTCCTCCCAAA |
| | i1b | CCTTGGCATTTACTGTGAAACTTG | CCTAAGTCAAGTACAACAATGCTGTG |
| | i9 | CGCTCTTGGCTAGCATCATACA | AGAGAAGGACCTGAGAGCAACTCT |
| | i19 | TGGATTTCCCAGCAGCTCTT | GCTTCCCAACCCCAACTTTT |
| | i39 | CAAAGATGGTCTTGAAATCTGTAGGA | AAACAGCTATTCTGGCACTGAAAA |
| | i77 | TTTGATTGAAGAGTAGGAAAGGTCAA | GGAAAGCCATTTGCACTGGTT |
| *CCSER1* | e1/e2 | CCCGGCCAGATTAATTTCTTG | TGATCGCCTGGATCCTGAGT |
| | e3/e4 | ACGGAAGCAGAGAGCAAGCT | CATCTAACTCACAGGAACCCAAGTT |
| | e5/e6 | CAAGGAAGAAAGAACAGATGCTAAAA | CCTTGTTCTACCTCCTTCGACAGT |
| | e7/e8 | GCACCACCTCACTCCCTGTT | CACTGAGCATTCATCTTTTATTCCA |
| | e9/e10 | TGTTTGCAGAAGGCTCTTGCT | GAGAATTTACATCATGGCCAAAAA |
| | i1a | TCCTTCCCCCTAGGTTATTTCTCT | TCAGTCCAACACCCTGAATGAG |
| | i1b | TGACGAGGTGGCCTTACATG | TCTGTGCCATTTTGTGACTT |
| | i5 | CTTCTTCAAGCAGGAGATTGTACTTG | CAAGCTGCTTTTGGCAGTGTAT |
| | i8 | GGGTTCCAAGTACTGCCATTCA | AACCAAAGCTCTGCTGCAAAC |
| *ACTB* | e1/e3 | CAGACATCAGGGTGTGATGGTTGG | GGGGTGTTGAAGGTCTCAAACATG |
| *FHIT* | e1/e2 | TCCCTCCCTCTGCCTTTCA | GATTTCTCTTCTTTCCTGAGCTTCA |
| | e2/e3 | TTGAAGCTCAGGAAAGAAGAG | TTCCACCGTCTGGATGTA |
| | e4/e5 | CTGAGGACTCCGAAGAGGTAGC | TCACAAGAGCGAAGGACAGTTC |
| | e6/e8 | TCTCACCTTTTCCATGCAGGAT | AACATGGACGTGAACGTGCTT |
| | e9/e10 | AGGAGGAAATGGCAGCAGAA | CAGGATCTGAAAAACATCTGTGTCA |
| *POLR2F* | e3/e5 | ATGTCGAGATCCTCCCCTCTG | GGCCTTGAGTTCCTTCATGG |
| *RPL11* | e2/e4 | AGCAGCCAAGGTGTTGGAG | TACTCCCGCACCTTTAGACC |
| *PPIB* | e4_1/2 | GTGAGCGCTTCCCCGATGAG | TGCCAAACACCACATGCTTGC |

**Supplementary Table 3. Primers used for quantification of mRNA levels and EU incorporation into nascent RNAs.** For *CCSER1*, manual alignment identified a different exon 10 from the one predicted by the Ensembl gene annotation system. RT-qPCR analysis confirmed that this exon is present in *CCSER1* mRNA in our cell lines (see Fig. 2a).

| Gene | Probe | PCR product | Forward primer | Reverse primer |
|---|---|---|---|---|
| *DMD* | Probe 1 (chr1:113,881,863-113,927,895) | 1_1 | TTCCCTATCTCTACCTGAAC | CACAATGACCCATGAATTCC |
| | | 1_2 | CCGATTTGTGTTTCAGTTTG | GATAATGTGACCCAACTCTG |
| | | 1_3 | TCTGGGCAAGACTAATTAAG | GATGGAGATCTGACCATAAC |
| | | 1_4 | TCTCCAACACAGGAAATAAC | GGCCTGTATTTACTTCAGTG |
| | | 1_5 | GTGAGATGATAGTGCAGTAG | CCCTAGTATATGGTGCTAAG |
| | | 1_6 | ATTAAGTCACAGCCATCAAC | ACAGGGTTCTTAAACACATC |
| | | 1_7 | TTTCAATTCCTTCGGGAATG | CAAGAGGTGCACTTGATTTC |
| | Probe 2 (chr1:114,902,820-114,948,473) | 2_1 | CTATCCAGCCAGTTCATTAC | GGACTGAGGACTAGGAAAT |
| | | 2_2 | TAAGGTGACTGGACTGTTTC | GCAGTGGAGATAATCCTTTG |
| | | 2_3 | GCTGTTGCTCATAATGATAG | TAACTGCTACAGGTAACATC |
| | | 2_4 | TTCTCAGTGTCTGTTGATAC | AACAAAGACCGAAAGTACAC |
| | | 2_5 | TTGCTTGGGTTTGTTAAAGG | AATTTGCTCATACGCACATC |
| | | 2_6 | GGAGGAGGACTACTTTAATG | GATTCTGTGTCAGACAAATG |
| | | 2_7 | TATGGACCAAACTCCTCTTC | GAAACTGTTTGCAGAATCAC |
| | | 2_8 | GCAGACTGTCTCAGTATAAC | AAGCCTAGGTAGTGATTAAG |
| | | 2_9 | CACGCATACATATACAAACG | AGGTGAAAGGAAGGTCTTAG |
| *CCSER1* | Probe 1 (chr4:34,954,291-34,999,441) | 1_1 | CCTCACCTCTATCTGACTAC | AGTGTCAACTCAGACCATAG |
| | | 1_2 | CAGATGGTTCTGTGTATTTG | TCTCACAGACCCGTTATTAC |
| | | 1_3 | GTGTTGCAATGCAGTAATTC | TCAAGCAGCTTTAGTAAAGG |
| | | 1_4 | TTTAGCATCTGTCCTTAGTG | GCAAGATATGGCAACTTATG |
| | | 1_5 | TAGTGCTGGCTTTGAATATC | TACAGCACAAATGGCAATAC |
| | | 1_6 | GCAGCAAGAAATAACTGATG | GCTTCAGAGGTGTATCTAAC |
| | Probe 2 (chr4:35,731,784-35,771,515) | 2_1 | CTACAGGAGCCTTAAGAATG | TTGACTAAAGGCAGGTAAGG |
| | | 2_2 | CTTTGCCCAGTTTAAGACAG | CATCCCTGCTATTATGAATG |
| | | 2_3 | CAGATTAGGTGATGTGATAG | TACTTTGAGGTAACGAGTAG |
| | | 2_4 | ATTGCACCTGTCAAACTAAC | TGCCAGTGATATTTAGAAGG |
| | | 2_5 | AAACACAGCTAAGTACTAGG | CTCATTTGGTGTTAACCAGA |
| *PARK2* | Probe 1 (chr3:43,706,102-43,758,694) | 1_1 | ACTCTGCTATTTCATCCTACTC | GATCCTGAGTTACTAACCTATG |
| | | 1_2 | CTTAATCCAACACCTAGAGATG | GCAGTTTACCTGATTCTTGTAG |
| | | 1_3 | TTCACTATAGATGGGCTCTATG | TATATGGAACTCGCTGTATCTG |
| | | 1_4 | CCCAACTCACTTTAGGTAATTC | GGTCCTGAGTATCTTCTAATTG |
| | | 1_5 | CTATACGCAAGACAATAGAGAC | CATCCCATGTCCTTCTAATTTG |
| | | 1_6 | TAATGGTACATCTGGAGGATAG | CTCTCATTGCTGAGTTAGAAAG |
| | Probe 2 (chr3:44,445,312-44,478,303) | 2_1 | ACTAAGTTACATGGGAGAAGTC | GAACAAAGAAAGAGGGAGAAAG |
| | | 2_2 | CATCTAGTGTCTCTTTCGTTAC | ATCAGTCAGTCTCTGGTAAATC |
| | | 2_3 | AGGGTGTATATTTCCCTCTATG | GGGTGCGGATATAATGTATTTC |
| | | 2_4 | GAGTGTTGCTAGTGGATATTTC | GAGGCAAATGAAGGATATAAGC |
| | | 2_5 | AGAATCCTATGGCTATACTCTG | GTAGCTAGCAAGTTCTACAATG |
| | | 2_6 | GCTACTTGTGTTGGATCTATTC | TGGAAACTCACTACCAGATAAG |

**Supplementary Table 4. Primers used to synthesize FISH probes and probe coordinates.** Coordinates are given according to the ICGSC/galGal4 chicken genome assembly.

| Cells | S-phase fraction | % of BrdU-labelled DNA in each fraction relative to total S-phase | | | Figure |
|---|---|---|---|---|---|
| | | Control early-S | Control mid-S | Control late-S | |
| WT | S1 | 38.84 | 6.00 | 13.38 | Figure 1e (exp. 1) |
| | S2 | 38.11 | 25.37 | 16.38 | |
| | S3 | 10.99 | 43.84 | 30.51 | |
| | S4 | 12.06 | 24.79 | 39.72 | |
| WT | S1 | 47.41 | 8.07 | 6.84 | Figure 2e (exp. 1) |
| | S2 | 42.47 | 29.28 | 7.89 | |
| | S3 | 7.06 | 45.63 | 25.39 | |
| | S4 | 3.05 | 17.01 | 59.88 | |
| WT | S1 | 30.71 | 6.00 | 3.17 | Figures 1e and 2e (exp. 2) |
| | S2 | 53.70 | 29.00 | 6.86 | |
| | S3 | 12.76 | 44.68 | 22.93 | |
| | S4 | 2.83 | 20.32 | 67.04 | |
| $DMD^{Tet/Tet}$ + tetracycline clone 1 | S1 | 42.46 | 4.73 | 3.29 | Figure 1f |
| | S2 | 43.10 | 31.37 | 12.27 | |
| | S3 | 10.12 | 43.79 | 35.23 | |
| | S4 | 4.32 | 20.11 | 49.21 | |
| $DMD^{Tet/Tet}$ + tetracycline clone 2 | S1 | 54.51 | 4.64 | 4.13 | |
| | S2 | 36.92 | 20.90 | 9.04 | |
| | S3 | 5.54 | 47.49 | 32.51 | |
| | S4 | 3.03 | 26.97 | 54.32 | |
| $CCSER1^{\beta a/\beta a}$ clone 1 | S1 | 36.30 | 3.66 | 4.32 | Figure 2f |
| | S2 | 47.20 | 30.74 | 9.66 | |
| | S3 | 12.89 | 51.43 | 32.74 | |
| | S4 | 3.61 | 14.16 | 53.27 | |
| $CCSER1^{\beta a/\beta a}$ clone 2 | S1 | 36.09 | 4.31 | 5.28 | |
| | S2 | 46.18 | 22.00 | 9.72 | |
| | S3 | 13.46 | 49.55 | 21.71 | |
| | S4 | 4.27 | 24.14 | 63.29 | |
| $DMD^{+/Tet}$ + tetracycline clone 1 | S1 | 42.43 | 3.06 | 2.63 | Figure 3c |
| | S2 | 45.11 | 24.55 | 11.01 | |
| | S3 | 8.39 | 50.55 | 38.50 | |
| | S4 | 4.07 | 21.84 | 47.86 | |
| $DMD^{+/Tet}$ + tetracycline clone 2 | S1 | 26.46 | 6.49 | 6.29 | |
| | S2 | 50.61 | 21.20 | 8.20 | |
| | S3 | 17.40 | 44.13 | 39.23 | |
| | S4 | 5.54 | 28.19 | 46.28 | |
| $DMD^{+/\beta a}$ clone 1 | S1 | 35.81 | 7.43 | 2.57 | Figure 3d |
| | S2 | 48.36 | 21.88 | 7.96 | |
| | S3 | 11.79 | 42.81 | 32.79 | |
| | S4 | 4.04 | 27.88 | 56.68 | |
| $DMD^{+/\beta a}$ clone 2 | S1 | 28.08 | 7.22 | 6.75 | |
| | S2 | 51.94 | 22.13 | 9.85 | |
| | S3 | 16.64 | 43.28 | 28.70 | |
| | S4 | 3.35 | 27.38 | 54.70 | |

**Supplementary Table 5. Replication timing control loci.** Quality control experiments confirmed enrichment of known early-, mid- and late-replicated loci in the expected fractions for replication timing profiles presented in Figs 1e, f, 2e, f and 3c, d.

| Gene | Name | Forward | Reverse |
|---|---|---|---|
| *DMD* | with(Tet) | CCTCCATAGAAGACACCGGG | TCGTCCCTGCAGATTGACAG |
| | with(βact) | CGTGCTGGTTATTGTGCTGT | CTGTCAATCTGCAGGGACGA |
| | without | CCCAGGAAAACATGTGGAGC | ATCTGTTGCAAATGAGATACAGCA |
| | both/p6.9kb | CGCTTCAGTTGCATTGGATGT | GGAATTGTTCACACGCTGGC |
| | p101.5kb | TGCTGCCGTTTTCTCAGATG | CACCAGGCCTGCTCTAAGCT |
| | p201.7kb | GGGTAGCTGTTCTTGGTTGC | ATGGTCCCTGGCCTTTTCAT |
| | p316.8kb | TGGATTTCCCAGCAGCTCTT | GCTTCCCAACCCCAACTTTT |
| | p391.4kb | CAAAGATGGTCTTGAAATCTGTAGGA | AAACAGCTATTCTGGCACTGAAAA |
| | p496kb | ATACCACATGACACCACGGC | AGCACCAATCAGCAATGCAA |
| | p598.2 | CTGGTTGTTTCTTTTGCGTGTC | ACATGGTGGGTGTCAAAGGA |
| | p701.1kb | GGCCTTTACCCGTCACCTTG | CAGTGCATGAGAAAGACCTGCA |
| | p798.8kb | CAAAGGAGAAGTGCAAGGCC | CAGCAAGGGAGAGTAGCTGT |
| | p899.1kb | AGTGTCTGCCCTTGTACCAC | TGCAAGCAGTTTTCCTCAGGT |
| | p995.5kb | GTCCATTGTGTGAACGGGTC | ATGGGGTTCCTACTGCAAGAA |
| *CCSER1* | p19.8kb | CACAGTGAAGACTGAGGGAAGGA | AATGAGGTGTAGCAAGATGCTCTCT |
| | p133.2kb | GCAGAACAGCCATAATTCAACTGT | TCAGTCTCAGCCTGATCTTCAAGT |
| | p217kb | CTTCTTCAAGCAGGAGATTGTACTTG | CAAGCTGCTTTTGGCAGTGTAT |
| | p318.9kb | CACAGGCTTTGACAGAATTGGA | CAGGTAGCAATGATCAAGGTACAGA |
| | p369.4kb | GGGTTCCAAGTACTGCCATTCA | AACCAAAGCTCTGCTGCAAAC |
| | p469.9kb | TGGCTGCAAATATAAGAATTGCTT | GCTCCAGAGAATGGTCACAACTG |
| | p571.6 | TGTTTGCTTGGAGGATGCAGTA | ATGGAACCAGCCATTTTTTGA |
| | p618.1kb | GCAAATACTACCTGACATGCAGCTT | AGCCTTCGGGTTCTGTAGCA |
| | p670kb | ATGGGTGTGTGTAAGTGGGG | CAAGGTAGTGCGTGAAATGGAA |
| | p769.8kb | GAAGCAAACAGACAGCGTGT | GCCAACCAAGAGCAAACCAT |
| Control regions | Mitochondria MIT | CATCCCATGCAAACTCCTG | GTAGTCCAGGCTTCACTTGA |
| | Early-replicated | GACGGTCAGGTTTGCCAAAG | TCCTGAGGATACGTTTTTCAG |
| | Mid-replicated | CAGGACAGGTATTCACA | GGCCTGAACACTGTGTCAAT |
| | Late-replicated | TGTACTTCTCTGTGGACATGCA | TGGCACAGAGGACAGGTAAGA |
| *FHIT* | i1 | TGTGCTGGGACCAATAGAAA | AACATCAGGTGCGAGTAGGG |
| | i3b | CATCCAGTGTCCAAGATGCATAC | CCCTGCCTCCCCTTCCT |
| | i3c | AGTGCATTGTCATTGAGTGATTTATG | GATAACCCCAAAGTGGACAAGTG |
| | i4b | GGCTTCGTGGGAGAAAAGG | AGATGCTGCTGTTGACAGTCGTA |
| | i5c | ACCGTAGGGAACCTCCTCTTG | GCTGCTCTTTGACCTTGAACAGT |
| | i5d | TCTCTCCTTTCAGGCTCCAAAT | AAGAAACAGGCCAAGTTAATGCA |
| | i7c | CAGCTCCCTTTCCTAGTCTGGAT | GGAAGCATTGATTGCAGATGAG |
| | i8b | TGGCGTGCACTTTCCTCTAA | CTGGTTTCCTTCTGATGCAACA |

**Supplementary Table 6. Primers used for replication timing analyses.**

| Gene | Motif | Probe | Forward | Reverse |
|---|---|---|---|---|
| *DMD* | A | A1 | ATAAACAGCACGAGAGTTGG | TTATCATGACTGCCAGAAAC |
| | | A2 | TTAATAGCGTGCTCAGGTTG | CCGGTGGAACTAACTGAAAG |
| | | A3 | TGAATTAAGCAGGCAGTTAC | AGTGCAATCAGTCTCATTTC |
| | | A4 | CCATATAGCCCTCCTATGTG | AACTGGTAACTTGTGTATCC |
| | | A5 | TGGTAAACTTGGGACTCATC | GGTGTGAAATTCCAGTAAAG |
| | I | I1 | TATTTATGGTACGGCAGTTG | CTCTGCACCAGAGGTATTTC |
| | | I2 | ACAGTCGAATGGAGGATATG | CCCAGCTTTCAAACCTTACC |
| | B | B1 | ATGCCTGATAGCAAATTCTG | CACCATCCCTCTTATTTCTC |
| | | B2 | TGACAAGTCTTGACCTAATC | TGAAGGGAAGGAATGAAGAC |
| | | B3 | CAATACATTCAGCTGTAAGG | GCTTATTTCACCCACTAATC |
| | | B4 | CCTCCCAGCTCAATATGAAG | CAAAGGCTAAATTGAGACTG |
| | | B5 | TTTGCTCTTGTCTCTTGATG | CATTAAACCCTCTGCCAATC |
| | | B6 | AAGCTTGGAAGCTACTAGTG | CACTTTAGTGTGGGATACTC |
| | | B7 | AGTAGAATCCCAGGTGAATG | AGCAAGTCTTCCTACCTTTC |
| | E | E1 | TGTGTTTGAGTGCTGATAAG | GAGAGGAAACTGGATGTTAG |
| | | E2 | GCATCCAGTTGTGTCTCTTC | GTGAACTCAATGCCATTTAC |
| | | E3 | ATCTCGCAGGAAGTTAAATG | GTGCCAGCAGAATATTAAAC |
| | | E4 | GGATGCCTAAAGTTGTACAG | TGCTCATGACTTGTACTATC |
| | | E5 | GCAGTGAGTATGAAGTCTTG | CTGTACCGGAGCAAGAAATG |
| | K | K1 | AGCAACTGCTTAAAGTAGTG | CAGGTGGTATATGACAAGAAC |
| | | K2 | CTAGAACCCTGGCACTTTAG | TACTAATGCGCTGACATGAC |
| | D | D1 | AATTACAGCCTTGCAGTACC | GTGCTAATACATGGAAACTC |
| | | D2 | GTGATAGATGCAGTTGTTTG | TTGATTGTCCCTAGAAAGTC |
| | | D3 | TTCACTGGTTCAAGATAAGG | CCAGAACATTTGGTGACTAC |
| | | D4 | CTGGTTTGGTTTGAGTAATC | TGCTTCTCTGTAGTACAATG |
| | | D5 | GTATACTGCACGTCAGCTAC | ATATCCCACTTCATGACTTG |
| | | D6 | TTGAATCCATGCCAGATGAG | TAGCTACACCTTGCTTTATC |
| | C | C1 | TGCTGAAGTGGAAAGTAAAG | AATATCAGTCCTGCCTTGTG |
| | | C2 | GACCTGAAGACATTCAGAAG | CAGTGATAGCCCTCTCAGAC |
| | | C3 | AGGAAAGGTATCTTGCTATG | GCCAAGAATAGTTCAATGAC |
| | | C4 | TGTATGTGAGTACACCTATG | GAGGGAGCAGACATTACTTG |
| | | C5 | GAGGCATTTAAGCTGGGAAG | GAAGGACAGATGAGCTAAAG |
| *CCSER1* | A | A1 | TTCCACCATCGTTGTTATGC | TGGAGGAAGTACGGTTACAC |
| | | A2 | GCACAGAGTGACATACTATC | AACACAGTCCTGGTTACAAG |
| | | A3 | AGGCGAGGTCTATGAGTTAC | CTCTTCCGAAGGAGCTTATG |
| | | A4 | GGGGAGGTAAATGTCCAATG | GATGGCATCTCTGACAATAC |
| | | A5 | TTTGAGTACCGGTCTACATC | CACAGAGCAGAACAGTTAAG |
| | D | D1 | TTCATGTAGGCATACGGAAC | CCCTTCTACCAGCATTAACG |
| | | D2 | GCGTTAATGCTGGTAGAAGG | TTCCATCCCACTGTTTACTC |
| | | D3 | TAGGTGCCTGACTAGTATGC | AGGTTGACAGTTCGAGTTAG |
| | B | B1 | GGGATTAGGAAGGCAATAGG | GTGACAGTTGGCATCGTTTC |
| | | B2 | GCTGGTAGTGTAACTGGTAG | CTACTTCTGGCATCGTATCC |
| | | B3 | ATCCTGTCACCTGCTAGAAG | CCTGAGGCACTTCGTGTATG |
| | | B4 | TGCCAGGGAATCATTGAAAG | AGCTGTGTACTGGAACAATC |
| | | B5 | TCAGCCTCCCTGAGAATATC | ACAGATGGCCAATTGTACTC |
| | | B6 | TGAGAGGCCATGTATGTTTC | TACTTTGTGTCTGCCTTGTG |
| | | B7 | GCAGACACAAAGTACGAAAG | TCAACATGTGGCTAGAAGAG |
| | | B8 | GCCACATGTTGAAATTCTCC | AAGGCAGAGCTATTTGAATG |
| | | B9 | TCATAGCAAAGGCCATGAAG | TAACCTAGCACTGTGAAGTC |
| | | B10 | CAGTGCTAGGTTAACAGTTG | GCATGAGTTCACCTATACAG |
| | | B11 | TAAACTGTGGTCTGCTCAAG | GGTCCTATCTCTGGCAATTC |
| | E | E1 | CAAAACTTGGGCCGATTTTG | TAAGCCTTGCCCTGAGATAC |
| | | E2 | AAGCGAATCTGTGGATATGG | GGCACTGGAAGCAATATTAG |

**Supplementary Table 7. Primers used to synthesize Morse-code probes.**

| Gene | Motif | Probe | Forward | Reverse |
| --- | --- | --- | --- | --- |
| *CCSER1* | C | C1 | GCCACCAGAAGTCATACAAG | CGTGCCAAAACCAATTAGTG |
| | | C2 | AAGAGGTGGGGCATTAATAC | TTCCCTTCTTGCAACTCTAC |
| | | C3 | GCTGCTAAGGGTTTGGTTAC | AACACCCACATGTACACATC |
| | | C4 | TGAAGGCAAGGATTGGATAC | ACAGAGAATAGCGGTTGATG |
| | | C5 | TCCTAAGAGCAACTTTCTTC | ATAGTTCCCGCTAAACCTTG |
| | | C6 | GTGCTTGGTACTACATTGTC | GCTTTTGAAGGCTTCTAGTC |

**Supplementary Table 7, continued.**
